# Free recall scaling laws and short-term memory effects in a latching attractor network

**DOI:** 10.1101/2020.12.19.423464

**Authors:** Vezha Boboeva, Alberto Pezzotta, Claudia Clopath

**Affiliations:** Department of Bioengineering, Imperial College London; The Francis Crick Institute

**Keywords:** free recall, episodic memory, recall capacity

## Abstract

Despite the complexity of human memory, paradigms like free recall have revealed robust qualitative and quantitative characteristics, such as power laws governing recall capacity. Although abstract random matrix models could explain such laws, the possibility of their implementation in large networks of interacting neurons has so far remained unexplored. We study an attractor network model of long-term memory endowed with firing rate adaptation and global inhibition. Under appropriate conditions, the transitioning behaviour of the network from memory to memory is constrained by limit cycles that prevent the network from recalling all memories, with scaling similar to what has been found in experiments. When the model is supplemented with a heteroassociative learning rule, complementing the standard autoassociative learning rule, as well as short-term synaptic facilitation, our model reproduces other key findings in the free recall literature, namely serial position effects, contiguity and forward asymmetry effects, as well as the semantic effects found to guide memory recall. The model is consistent with a broad series of manipulations aimed at gaining a better understanding of the variables that affect recall, such as the role of rehearsal, presentation rates and (continuous/end-of-list) distractor conditions. We predict that recall capacity may be increased with the addition of small amounts of noise, for example in the form of weak random stimuli during recall. Moreover, we predict that although the statistics of the encoded memories has a strong effect on the recall capacity, the power laws governing recall capacity may still be expected to hold.

## 1 Introduction

Despite the impressive capacity of the human brain to store information, its ability to recall it can be unreliable and fraught with mistakes. Many different task designs have aimed at quantifying this unreliability. In particular, in the free-recall task, participants are sequentially presented with a list of unrelated items. Immediately after a cue signaling the end of the list, they are asked to recall these items in any order. The absence of experimenter-imposed structure provides relatively unconstrained conditions expected to reveal insights into the natural memory processes guiding recall, such as the interaction between short-term and long-term processes. One of the most salient findings in the free recall literature is the limited recall capacity of human subjects. With an increasing list length, participants recall an always diminishing fraction of the presented items. Studies have reported that this follows a power law in the length of the list [1–6]. But how does this limited recall capacity arise, when instead the long-term storage seems virtually unlimited? Studies into attractor models of memory have shown that within the storage capacity limit, all memories, theoretically, should be retrievable, when fast noise levels are sufficiently low [7–10]. This raises the possibility that additional mechanisms hinder the retrieval of memories, but it is unclear how such scaling laws emerge from large networks of interacting neurons.

This limited recall has many characteristics that have been extensively studied, stimulating much of the theoretical discussion regarding the memory processes underlying free recall. One benchmark finding pertains to serial position. It has been found that recall is excellent for the last items presented, less accurate although remarkably good for the first items in the list, and poor for the middle list items. These features, known as the primacy and recency effects, respectively, are robust findings that have been replicated many times, and extend to many other memory paradigms [11]. Traditionally, the U-shaped serial position curve has been explained by invoking two distinct but complementary mechanisms [12]. In this view, the primacy effect reflects the advantage in processing (e.g. in terms of the number of rehearsals) given to the first items in the list, resulting in the selective transfer of the early items into a stable long-term memory store). By contrast, the recency effect would be interpreted as the output of a temporary and highly accessible short-term store of limited capacity.

The primacy effect has been linked to selective rehearsal in several different manipulations of the basic paradigm. When participants were encouraged to rehearse aloud the item of their choice during the presentation of the words, the first items in the list received far more rehearsals than the middle and last items [13]. Conversely, when participants were instructed to rehearse only the item that was currently being presented, the number of rehearsals was equated across the stimuli, and the primacy effect was greatly attenuated [14, 15]. Furthermore, the primacy effect is reduced when free recall is performed at a fast presentation rate [16, 17], with concurrent articulatory suppression paradigms [18], or with paradigms utilising a filled interval between each item in the list [19–21], and under incidental learning conditions [22, 21, 20, 23], providing behavioural evidence for the rehearsal hypothesis. Also, the magnitude of the primacy effect seems to decrease the longer the list [24].

The recency effect, instead, has long been thought to be the result of the output of a fragile short-term store. It has been found that a period of distractor activity immediately after the end of the list and preceding recall eliminates the recency effect but does not affect performance on earlier list items (e.g., [25]). Interestingly though, recency effects are still observed when the distractor is placed into the interstimulus as well as at the end of the list, posing a problem for the short-term store interpretation [26].

Recency effects are observed not only in terms of the global position of items in the list, but also in terms of their position relative to other items. These are highlighted by *transitions* that participants make when recalling items. In other words, participants are more likely to recall items that are close in the list consequently, rather than items that are far apart in the list consequently, even if they are free to choose to recall in whatever order that they wish. Moreover, they are more likely to recall items in the order they were presented in the list (forward contiguity) rather than in the reverse order (backward contiguity). This finding, called the lag-recency effect, has been replicated under many different experimental manipulations, and persists under all of the manipulations described so far [26].

However, the formation and use of associations between words due to their temporal proximity in the list is only one element guiding memory search. When a pair of semantically related words is included in the presented list, the related words are often recalled successively [27]. Evidently, measuring such effects requires making assumptions about the semantic relatedness of words, and a variety of standardised similarity metrics have been designed and used in order to systematically quantify the meanings of words [28, 29]. Still, such effects have been found to be robust against the specific metrics used to measure semantic relatedness, and suggested to be a universal search strategy in human subjects [30].

Here, we propose a network model of free recall in which both temporal and semantic cues guide memory search. We do not make any distinction between the long-term and the short-term store, and we consider that both are subserved by the same memory representations in the model network. What distinguishes their different properties are the plasticity rules that govern dynamics at different time-scales. Under appropriate conditions, the network learns and endogenously reactivates memories even when the corresponding external stimuli are not present. The recall capacity of our network is a power law of the list length. Importantly, this limited recall capacity is not a consequence of limited storage capacity, that is much higher for the network. Rather, it can be traced to the emergence of limit cycles that trap the network’s ability to explore memory space. These limit cycles arise as a result of correlations between memory patterns [31], without which the network is found to explore pattern space more homogeneously. Indeed, we find that when memory patterns are orthogonal to one another, the recall capacity is not limited.

In order to better understand the dynamics of the spontaneous reactivations, and the conditions under which they arise, we develop a mean-field theory of our model under several simplifying assumptions (see Supplementary Material). Similar to other models [32–34], we find that the addition of firing rate adaptation and global inhibition generates additional phases of operation to the default “ferromagnetic” and “paramagnetic” regimes that have been extensively studied [35–37, 9]. We derive the conditions under which these additional phases, characterised by an oscillatory mode of reactivation in the single memory storage scenario, and random reactivations when multiple memories are stored, arise. This latter phase, the latching phase [38–41], is the regime of interest in which our model operates.

Our model reproduces several benchmark findings in the free recall literature, such as serial recall positions, temporal contiguity, asymmetry effects and displays a similar pattern as some of the manipulations of the basic paradigm, aimed at controlling rehearsals. Furthermore it reproduces some of the semantic effects guiding memory search. Our model displays recall capacity that is a power function of the list length, as has been found in experiments and suggests a strong role for both temporal and semantic correlations in guiding the recall trajectory. We predict that recall capacity can be increased with the addition of small amounts of noise, for example in the form of weak random stimuli, or weak distractors into the dynamics, and that the structure of the encoded memories has a strong effect on the recall capacity.

## 2 Results

### 2.1 Limit cycles lead to limited recall capacity

It is well known that human subjects’ ability to recall items from memory, in the absence of external cues, is limited [1–6]. To test the possible underlying mechanisms of limited recall capacity, we build a model which initially has stored *p* long-term memories through a prescription (Eq.(1)). We then present the network with a set of *L* memories sequentially, out of a total of *p* long-term memories in the form of input (*I*_cue_) to specific neurons coding for that memory, with pauses in between items (*T_ISI_*) (Fig. 1A and B). The stimuli lead to a strengthening of the weights (Fig. 1 C) through a Hebbian term composed of both autoassociative (co-activation leads to potentiation) and heteroassociative components [42–44] (see Methods, Eq.(12) and (13)), as well as short-term facilitation. During the pauses in between each presented item, the network does not sustain the stimulus, and the stimulus-specific activity decays to zero. This is because the network is equipped with neuron-specific rate adaptation and global inhibition. However, the network spontaneously reactivates (see Supplementary Material, and Fig. S1) and retrieves a random memory item, mostly among those that have been presented so far (Fig. 1 B). In addition, adaptation prevents the network from retrieving the same item at intervals shorter than the adaptation time scale.

**Figure 1:**
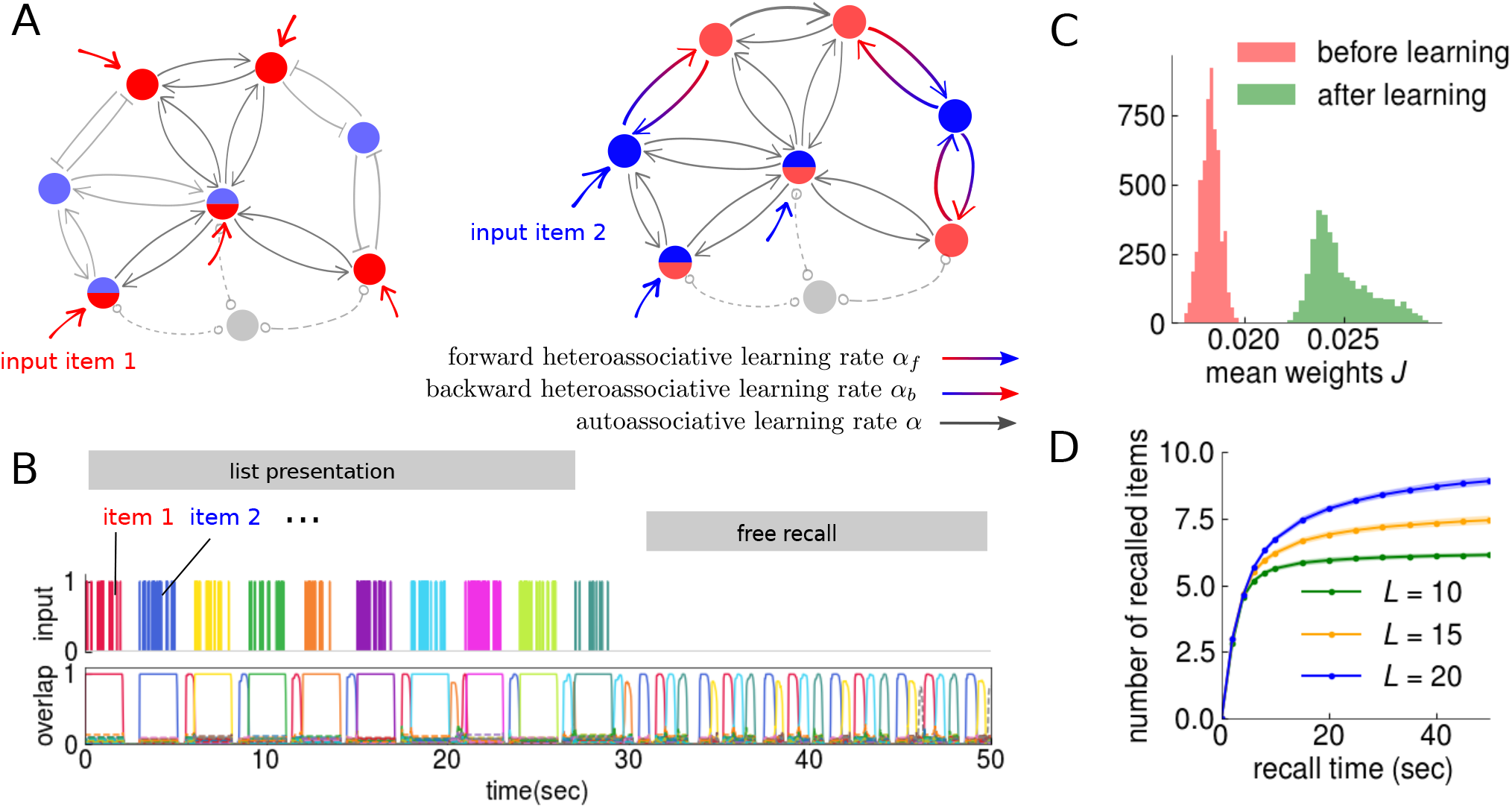
Limit cycles lead to limited recall capacity. **(A)** We consider a model of long-term memory [45] equipped with dynamic auto- and heteroassociative weights, in addition to the baseline weights (details in Methods). *a* is the autoassociative learning rate, whereas *α_f_* and *α_b_* are the forward and backward heteroassociative learning rates, shown in coloured arrows with colour gradients. Coloured arrows correspond to input to specific neurons coding for a given memory item (each colour corresponds to a memory item). Gray arrows correspond to the baseline weights. **(B)** We simulate our model with a free-recall task: the presentation of an item corresponds to the addition of an external input field to all the neurons coding for that item (red and blue arrows in (A)); items are presented with a pause in between (*inter-stimulus interval*, *T_ISI_*); after the presentation of the list, the network is let to freely recall items. Recall is measured through the overlap, or the correlation between the network state and a given memory, as a function of time (the different colours correspond to different memories). **(C)** The autoassociative learning encodes individual memory items: the distribution of the mean weights, *J*, between neurons belonging to a given memory can be seen to increase its mean and standard deviation after list learning. **(D)** The recall capacity of the network is limited, as expressed by the plateau in the number of recalled items. This limited recall occurs for different values of the list length (different colours).

At the end of the stimulation, the external input is shut down in order to test the ability of the network to recall the list. If the learning rates (both auto and heteroassociative) are set to be too low, the network’s activity either dies out after recalling a few items, or performs excursions to memories that have not been presented in the list (Fig. S4 bottom). If, instead the learning rates are high enough, the network’s activity is constrained to the items in the list, performing recall (Fig. S4 top). However, although the network almost always recalls items from the list, this recall is limited, and the number of items recalled soon reaches a plateau (Fig. 1 D). This limited capacity can be seen to arise from the limit cycles the network enters (Fig. 1 B). Altogether, our model suggests that sufficiently high learning rates and firing rate adaptation can lead to spontaneous recall dynamics that is largely limited to the list, albeit still limited to a fraction of the list, as in free recall experiments [1–6].

### 2.2 Reactivations and short-term plasticity lead to serial position effects

Human subjects display a variety of biases when they are asked to freely recall list items. Some of these biases relate to the position of the item in the list: recall is usually good for the last and first items, but poor for items in the middle of the list. These are robust effects that have been found in a variety of paradigms, including free recall, but the mechanisms through which they arise are still not well understood.

Many manipulations have aimed at uncovering the factors that affect recall. One of these consists of varying the presentation rate. It has been found that increasing the presentation rate negatively impacts the recall of the first items in the list, but does not markedly affect the last items in the list. We sought to determine whether our model could reproduce this finding. We varied the interstimulus interval, *T_ISI_*, thereby changing the presentation rate, and found that the recall for the initial items was poorer to the extent that *T_ISI_* is lower (Fig.2A). This can be understood by analysing the differential roles of the long-term weights *J* and the synaptic facilitation *u*. While the long-term weights become stronger for the first items, due to more frequent reactivations; the short-term presynaptic facilitation, instead, decays to baseline over the short-term plasticity timescale (Fig.2 B). Under the default ISI, the combined effect of the two is an effective synaptic strength curve *Ju* with its characteristic non-monotonic U shape (Fig.2C). Instead, under short ISI, *J* is more flat and therefore the U shape disappears (Fig.2B, red curve).

**Figure 2:**
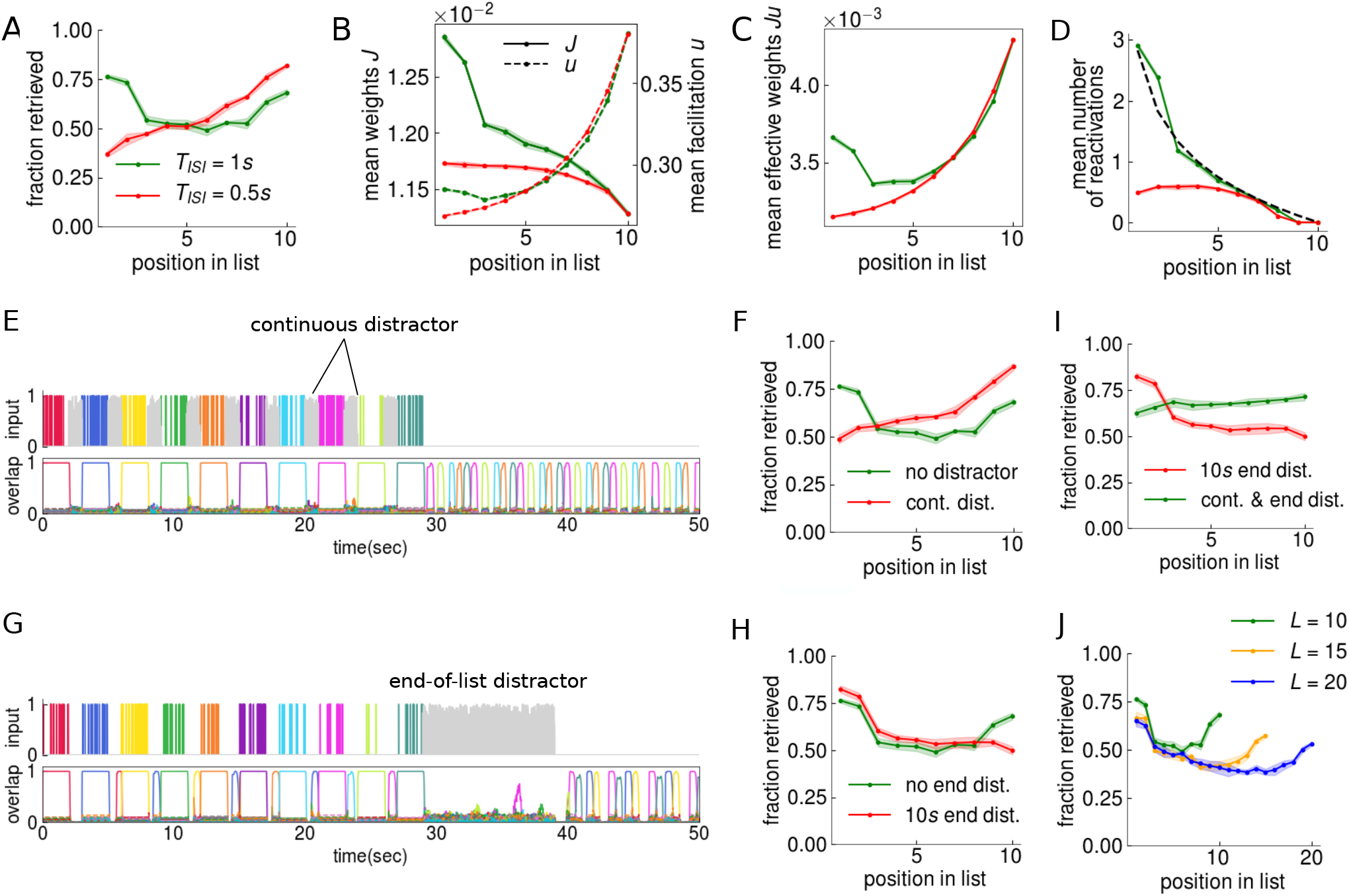
Reactivations and short-term synaptic plasticity lead to serial position effects. **(A)** Effects of the interstimulus interval (ISI) on recall are explored. Reducing the ISI impairs the primacy portion of the serial position curve, while leaving the recency portion intact. **(B)** This can be traced back to the mean weights (*J*) and synaptic facilitation (*u*) of the memory items in various positions. When the ISI is high enough (green curve), the first items benefit from more frequent reactivations during the ISI, leading to a strengthening of the weights for items at initial positions. By contrast, the last items benefit from synaptic facilitation that has not yet decayed to baseline. When the ISI is reduced (red curve), reactivations occur less frequently; as a result, the weights for the initial items are not strengthened. The synaptic facilitation, instead, is less affected. **(C)** The combined effect of both weight changes (*J*) and synaptic facilitation (*u*) lead to both primacy and recency effects, when the ISI is sufficient (green curve). **(D)** The weight changes can be traced to the network reactivations. When the presentation rates are slower (green curve), reactivations occur roughly once per pause, randomly with equal probability 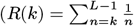, plotted in black). When the presentation rates are faster (red curve), the network’s ability to reactivate items is reduced, and with it the primacy effect, as shown in (A). **(E)** We model distractors by perturbing our network during the pauses with random input to all neurons (gray). As a result, the network cannot spontaneously recall memories (different colors). **(F)** As a result, as with reducing the ISI, the continuous distractor eliminates the primacy effect (red curve), through the elimination of reactivations. **(G)** Distractors can also be placed after list presentation. **(H)** In this case, the recency is eliminated (red curve) compared to the basic paradigm. **(I)** When both manipulations (continuous and end-of-list) are paired, however, the recency effect is mildly restored, as has been reported experimentally. **(J)** Serial position effects persist also when varying the list length *L*.

It was hypothesised early on that human participants may have been rehearsing earlier items more frequently, thereby consolidating their memory of it. Consequently, the overt rehearsal paradigm was designed, in which participants were asked to say aloud whatever they had been rehearsing silently. It was found that participants tend to rehearse earlier items more frequently [46]. Given that spontaneous reactivations are a feature of our model, and that memory items are presented to the network sequentially, our model naturally reproduces a larger number of reactivations for the first items, when the interstimulus interval is large enough (Fig.2D).

Another simple manipulation that negatively affects recall of the first items in the list is the continuous distractor paradigm, in which participants are asked to perform unrelated tasks (such as simple additions) in between pauses between list items [19–21]. We reasoned that if reactivations are necessary for the recall of the first items, then filling the interstimulus intervals, just as reducing the *T_ISI_*, must also compromise the primacy effect. We tested this idea by filling the pauses with a uniformly distributed random input to the whole network (shown in gray, in Fig.2E). Similarly to the previous manipulation, the ability of the network to spontaneously reactivate the items is again compromised (see also Fig. S3 D), and with it the primacy effect (Fig.2F).

Conversely, none of the two previously mentioned manipulations affect recall of the last items in human participants. Delayed recall, however, in which participants are asked to perform an unrelated task for a given duration after the list, does affect the recency portion of the curve, to the extent that this duration is longer [25]. We sought to verify whether our model was consistent with this finding. While the first two manipulations affecting reactivations did not affect recency (Fig.2A and Fig.2F), extending the duration after the list is presented and filling it with random input (Fig.2G) compromises the recall of the last memory items (Fig.2H).

Interestingly, although delayed recall impairs the recall of the last items, when it is paired with the continuous distractor paradigm in which rehearsal is minimised, the recency effect is restored. This finding and its subsequent replications under a variety of experimental conditions [19, 47] has remained problematic for dual-store models (at least in their most simple variants) [26] and the ensuing attempt to develop single-store models of memory that account for recency across various time scales. We also applied this manipulation to our model and found a similar behaviour in our network, although this effect was mild; a period of distractor activity at the end of the list eliminates the recency effect, but when it is paired with the continuous distractors, recency is somewhat restored (Fig.2I). This is because despite the manipulation, the effective weights *Ju* of the last items are still higher than those of the first items. Finally, we find that our model reproduces both primacy and recency effects under increasing list length, as has been found in experiments [24] (Fig.2 J).

### 2.3 Time-dependent heteroassociative learning leads to temporal contiguity of recall

Although serial position curves have provided much insight into memory processes, particularly in relation to the debate between short and long-term stores, they do not provide much information about the dynamics and the mechanisms that guide memory search. One such mechanism relates to how the temporal succession of items contributes to their subsequent recall. To this end, the conditional response probability measure (CRP)was designed [48]. The CRP, measured as a function of the lag, expresses the probability that the next recalled item is presented in position *i +* lag, given that the current recalled item is presented in position *i*. The CRP expresses the finding that subjects are more likely to recall successive items in the order they were presented in the list, findings known as the *contiguity* and the *forward asymmetry* effects. The lag dependence of the CRP is a robust property of free recall [48], and persists under a variety of experimental manipulations, such as presentation modality (visual vs. auditory), list length, and presentation rate. They also persist in delayed as well as continuous distractor conditions, designed to minimise rehearsal [26].

We reasoned that the heteroassociative component of the long-term learning rule [42], may lead to contiguity in the recall sequences uttered by our network. Consistent with this idea, we found both contiguity effects as well as the forward asymmetry effects characteristic of free recall (Fig. 3A). The forward contiguity effect is achieved through the weights linking subsequent memory items in forward order, that is, from the presynaptic neurons of item *i* to the postsynaptic neurons of item *i* + 1. This translates into a block diagonal weight matrix with positive values not only within the blocks (coding for single items), but also in the upper triangle (Fig. 3B). Equally present but to a lesser extent, the backward contiguity effect is similarly achieved through the strengthening of the weights between presynaptic neurons of item *i* + 1 and the postsynaptic neurons of item *i*, as can be seen in the lower triangle of the weights matrix. The asymmetry is achieved through the differential magnitudes of the forward and backward learning rates (see Eqs.(12) and (13)) used in the updating of the ingoing and outgoing weights of each neuron (see also Fig. S5).

**Figure 3:**
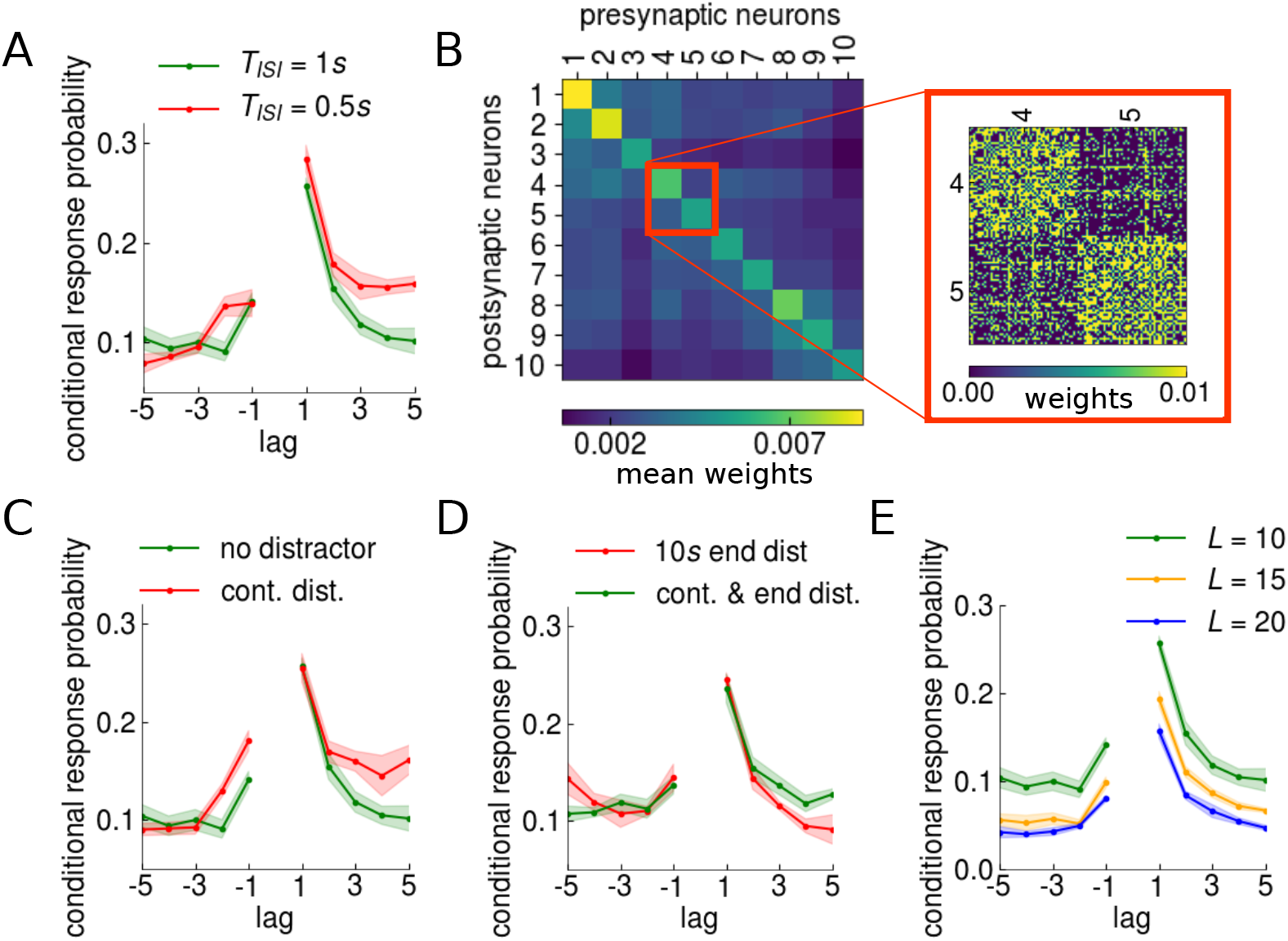
Time-dependent heteroassociative learning leads to temporal contiguity of recall. Experimentally observed properties of recall which depend on relative positions of items in the list are captured by our model. **(A)** The conditional response probability (CRP), measured as a function of the lag, expresses the probability that the next recalled item is presented in position *i* + lag, given that the current recalled item is presented in position *i*. Changing the interstimulus interval duration, *T_ISI_*, has a small effect on the lag-CRP: shorter ISI lead to a higher forward portion of the curve (red curve) whereas longer ISI to a lower forward portion of the curve (green curve). **(B)** Higher probabilities to make transitions with small lags (temporal contiguity) can be understood through the difference between the weights matrix before and after list presentation. When the weights between neurons coding for list items are plotted in order of presentation (*i*_1_ to *i*_10_), the higher values of the forward weights (upper triangle), relative to the backward weights (lower triangle), become apparent. This is especially apparent in the right panel of the figure, in the zoom of the weights between items *i*_2_ and *i*_3_. This effect is achieved through the heteroassociative learning rule with a higher forward learning rate *α_f_*, relative to the backward learning rate *α_b_* (see also Fig. S5). **(C)** When a distractor fills the ISI (red curve), the backward portion of the lag-CRP becomes slightly higher, relative to no distractor (green curve). Still, the overall CRP curve is robust to this perturbation, as has been found in experiments. **(D)** A period of end-of-list distractor (red curve) or continuous distractor coupled with end-of-list distractor (green curve) does not markedly affect the CRP. **(E)** Lag-CRP measures persist also with increasing list length.

The temporal proximity of items due to fast presentation rates, may lead to the formation of stronger inter-item associations. However, even longer presentation times, allowing for extensive rehearsal have been found to produce salient contiguity and asymmetry effects [48]. The continuous distractor condition, instead, may be argued to disrupt such inter-item associations between presented items from taking place. We tested both manipulations in our model, varying inter-stimulus intervals, and the continuous distractor, and we found that although the magnitudes of the contiguity and the asymmetry are indeed affected, the general trend is unaffected (Fig. 3 A and C). In our model, decreasing the inter-stimulus interval duration increases the magnitude of the forward lag-CRP. This can be understood as being due to the activity trace of the previous item that has not fully decayed to baseline, when the current item is presented. When inter-stimulus intervals are filled with distractor activity, instead, a different mechanism, mainly the absence of reactivations, allows for stronger backward associations to take place. Pairing both of these manipulations, distractor between the stimuli, and distractor at the end of the list, does not markedly change the lag-CRP (Fig.3D).

In addition to the increased probability to recall items in close proximity, it has been found that participants sometimes recall items further apart in a consecutive manner. For instance, the majority of participants tend to initiate recall by one of the last items in the list [30], after which they recall one of the initial items. In our model, this heightened probability to make large jumps backwards is a byproduct of the backward heteroassociative component of the learning rule, coupled with the spontaneous reactivations: the weights from the presynaptic neurons of a given presented item to the postsynaptic neurons of the previously reactivated item are strengthened. Given that earlier items have more possibilities to become reactivated, this increases the probability of large jumps backwards, as expressed through the non-monotonic backwards portion of the lag-CRP curves (Fig. S5).

Finally, we find that although the magnitudes of the lag-CRP are reduced with increasing list length, they still persist (Fig.3E), consistent with what has been found experimentally [48].

### 2.4 Transitions between recalled items are guided by correlations between memories

It is well known that participants do not rely solely on temporal cues to guide memory search, they also make use of existing semantic or associative relations between list items [49, 27]. A distinct line of computational research, on Potts models of long-term memory based on distinct short and long range connections, has shown the tendency of the network to make transitions between memories, a type of dynamics called latching dynamics. Such a process is to be attributed to firing rate adaptation and local and global inhibition, under the influence of which memory attractors are perpetually destabilised [32–34, 38–41]. It has been shown that such transitions usually occur between memories that are correlated with one another [39–41, 50, 51]. Since firing rate adaptation and inhibition are both features of our model, we reasoned that such correlations between memory representations may serve as cues for “semantic” transitions, distinct from the temporal cues giving rise to the contiguity effect. We tested whether such recall by semantic associations was present in our model by applying the semantic conditional response probability measure to our recall sequences. The semantic CRP measure is basically a generalisation of the lag-CRP measure from the temporal dimension to the semantic dimension [52]. In our case, we consider the semantic similarity to be expressed in the correlation between the memory items, computed as the dot product between their representations. We found a strong positive correlation between the correlation between list items and the semantic-CRP (Fig. 4 A).

**Figure 4:**
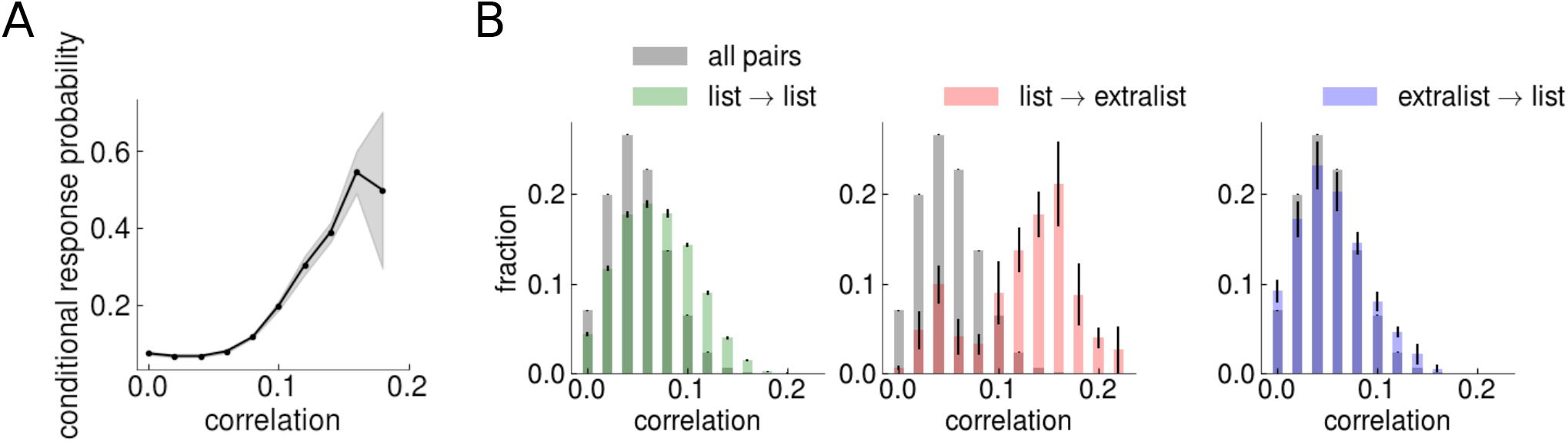
Transitions between recalled items are guided by correlations between memories. **(A)** The semantic-conditional response probability expresses the tendency of the network to recall pairs of items with higher correlation consecutively. The correlation between two memory items is calculated simply as the dot product of their representations. Note that for even higher correlations, the CRP decreases because pairs of items with such large correlations become increasingly rare. **(B)** The distribution of correlations between memory patterns involved in a transition differentiates among intra- and extra-list items. All distributions have been normalised. Left: black corresponds to the distributions of the correlation between all pairs of memories stored in long-term memory (black), and green to pairs in the list among which at least one transition has occured (green). The distribution of list → list transitions, has a slightly larger mean and standard deviation. Middle: the number of items encoded in the network can be much larger than the number of items presented to the network in the list, and occasionally one of these extra-list items is erroneously recalled. Pairs of items, list → extralist, between which at least one transition has occurred, shows a significantly larger mean and standard deviation, suggesting that large correlations may drive mistakes in recall. Right: the distribution of transitions from extra-list items back to intra-list items, however, displays a similar mean and standard deviation to that between all memory items, as these are driven by stronger weights for intralist items, a result of list learning.

A different, but related finding pertains to recall mistakes: transitions to such intrusions usually occur when there is a semantic relation or association between the intrusion and an item in the list [53]. Such intrusions are naturally expected to occur in our model, as we assume that a large set of memories has been previously stored in its synaptic weights (Eq.(1)). Occasionally, extra-list items are incorrectly recalled in our network. We tested whether such intrusions occur due to stronger associations by measuring the correlation between two types of latching transitions: transitions to items present in the list as well as transitions to items not present in the list. We found that on average, items belonging to the set of transitions list→ extra-list have a higher pairwise correlation than the set of items belonging to the set of transitions list→ list (Fig.4B), suggesting that when intrusions do occur, they do so because of the higher correlation they have with the previous item. Taken together, our results suggest a strong role for semantic associations in memory search in our model, affecting both correct as well as false recall, even though the items in our model are only randomly correlated.

### 2.5 Recall capacity is a power law of the list length. Small amounts of neuronal noise increases recall capacity

In the previous sections, we have seen that both temporal and semantic correlations in our model play an important role in guiding the transitions occurring between memory items. Interestingly, such mechanisms have been proposed to explain another basic finding in the free recall literature: power laws governing recall capacity. In this class of abstract models, the recall trajectory is completely deterministic, and depends on the similarity between items [31, 54]. By making the assumption that the recall trajectory is guided by the highest correlation, scaling laws similar to that of experiments have been derived [6]. This is due to the fact that the network dynamics eventually enters a cycle, repeatedly recalling the memories it has already visited, and no more new items can be recalled. This predicts that recall terminates when the same items start to be repeatedly retrieved, a hypothesis that has been previously proposed [55]. In support of this idea, human subjects usually recall items once, but when they report the same item for the second time, there is a higher likelihood that no new items will be recalled [56]. The scaling laws are general properties of sparse random graphs (in this case the correlation matrix), and it has been hypothesised that they may be independent of the particular mechanism generating the transitions between the items. We test this hypothesis in our model, in which such transitions occur because of the destabilising effect of firing rate adaptation. Our model stores a large number of sparse, long-term memories through a Hebbian prescription rule [45] (Fig. 5 A, Eq.(1)). For simplicity, we do not explicitly simulate a learning phase here (see Eq.(2) in Methods). Initially, we cue the network with an external input to neurons coding for a memory. Under appropriate conditions (see Supplementary Material), the network continuously recalls memories, even when external input is removed (Fig. 5B, top panel). We collect statistics until steady state is reached, which is characterised by a plateau in the fraction of items retrieved (Fig. 5 C). This plateau is due to the appearance of limit cycles (Fig. 5 B). We find that the number of recalled items is a power law of the number of items in the list (Fig. 5D).

**Figure 5:**
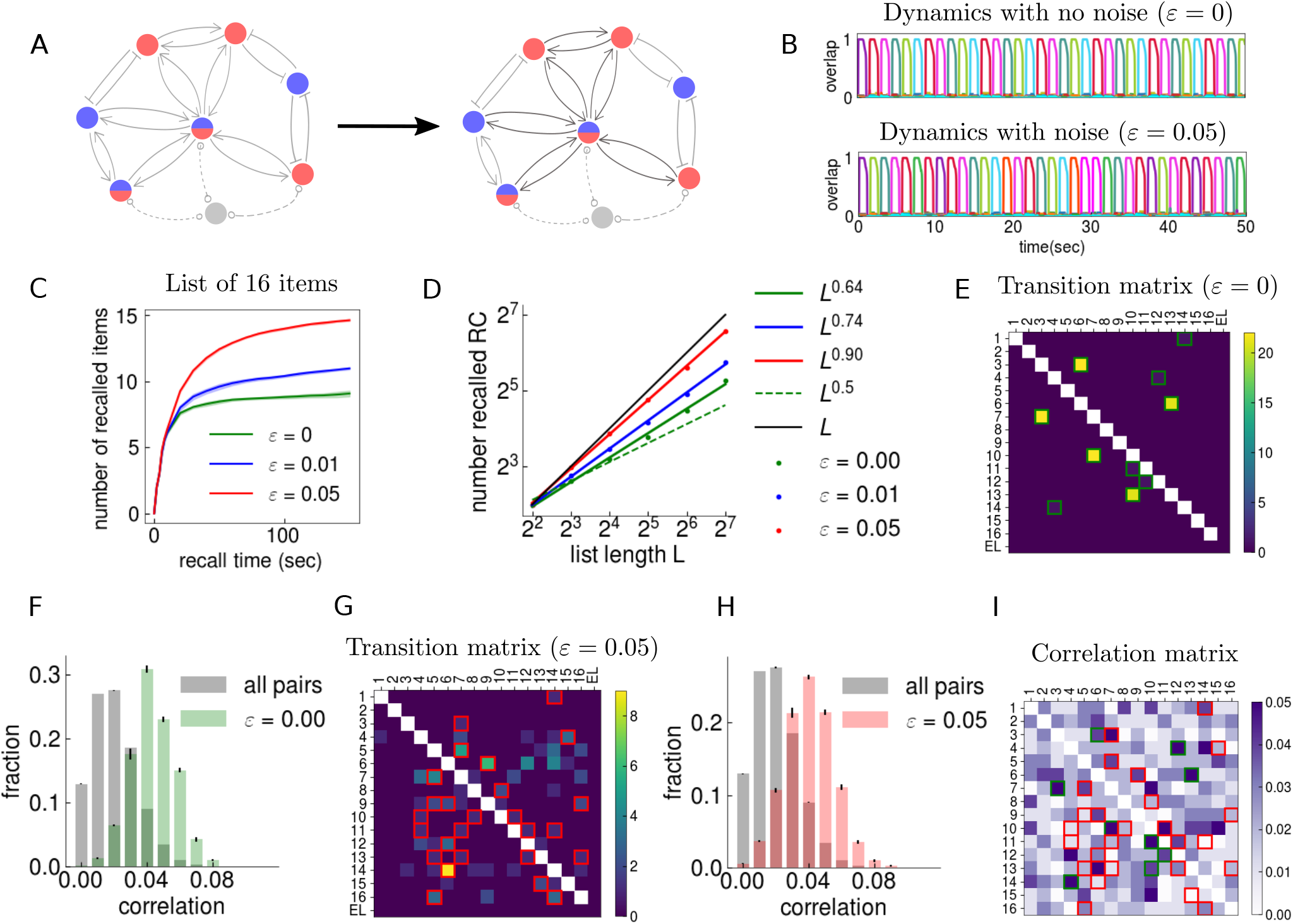
Recall capacity is a power law of the list length. Small amounts of neuronal noise increase recall capacity. **(A)** We study an attractor neural network that stores a large number of sparse, long-term memories through a Hebbian prescription rule [45]. We model list learning through the enhancement of the weights (shown in light/dark gray) coding for a subset of the memories (shown in red and blue), and we allow for firing rate adaptation and global inhibition. We also explore the effects of a noise term (*ε*) to the input of each neuron. **(B)** Our network displays latching dynamics, hopping from memory to memory. This is shown through the overlap, or the correlation between the network state and a given memory, as a function of time (the different colours correspond to different memories). However, the ability of the network to recall all of the items is restricted by limit cycles. Top: no noise *ε* = 0. Bottom: noise *ε* = 0.05 (corresponding to 5% of signal from the recurrent weights). **(C)** As a result, the fraction of recalled items saturates to a value less than unity in the long time limit, increasing with noise (corresponding to the different coloured curves). **(D)** The number of recalled items scales as a power law of the number of items. Dots correspond to simulations with different noise levels (different colours) while lines are power law fits (exponents reported in the legend). The black line is perfect recall *RC* = *L*, whereas the green dashed line is the quantity 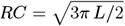, recently found to provide a fit to human free recall data [6]. **(E)** The transition matrix for one simulation without noise, where entry (row, column) = (*i, j*) corresponds to the transition from memory item *i* to memory item *j*. The last column and row correspond to transitions to and from extra-list memory items (EL). The colourbar expresses the number of times the transition has occurred. The network only makes certain transitions. For each row, the column corresponding to the most frequent transition (if any) is highlighted in green. **(F)** The transitions between memories are guided by large correlations, although not deterministically, and low-correlation transitions also occur. The gray distribution corresponds to the pairwise correlation values between all memories stored in the network, while the green distribution corresponds to the correlation values between pairs between which at least one transition has occurred, in the noiseless network. **(G)** The transition matrix when noise (*ε* = 0.05) is present. The addition of noise into the dynamics allows for the network to recall more items, increasing the recall capacity. For each row, the column corresponding to the most frequent transition (if any) is highlighted in red. **(H)** Same as in (F), but for the network with noise *ε =* 0.05. The addition of noise allows for more lower-correlation transitions to occur, as expressed by the lower mean of the red distribution, relative to the green one in (F). **(I)** The correlation matrix corresponding to the items presented in (E) and (F). The green squares highlight the most frequent transitions in the absence of noise (highlighted also in panel (E)), whereas the red squares highlight transitions when noise is present (highlighted in panel (G)).

The addition of some noise into the input of every neuron (*ε* = 0.01 – 0.05, corresponding to 1 – 5% of signal from the recurrent weights) induces some stochasticity in the recall sequences (Fig. 5B, bottom panel), and increases the exponent of the power law, eventually allowing for all of the list memory items to be recalled (Fig. 5 D). We find that in the noiseless network, transitions between memories occur largely deterministically (Fig. 5 E), but they do not always occur between the highest correlation pairs, even though they do occur between higher-correlation pairs (Fig. 5 F). When noise is present, transitions become more stochastic (Fig. 5 G) and occur between lower-correlation pairs (Fig. 5H, I). Note however, that the amounts of noise we have considered here are very small. A large amount of noise, instead, shuts down all activity in the network and hinders the recall of any memory (see Supplementary Material and Fig. S3 D). We have used this to model distractors in the full model (see also Fig. 2E and G).

### 2.6 The structure of the stored memory set exerts a strong influence on the recall capacity

Given that in our model, correlations between memories largely drive transitions during recall, we reasoned that completely eliminating them should allow the network to explore all possible transitions. We tested this possibility by simulating the network storing orthogonal patterns. We found that with orthogonal patterns, the network does not enter into limit cycles (Fig. 6 A), and therefore the recall capacity is not restricted, and is equal to the number of items in the list (Fig. 6B). In this case, though, the network takes more time to reach the maximal recall capacity, due to bouts in the dynamics in which it goes back and forth between the same pair of memories (Fig. 6 A), as can also be seen in the transition matrix (Fig. 6 C). However, a direct comparison of how the recall capacity scales with the list length *L* for the same set of parameters as the model with random patterns was not possible due to the limited number of orthogonal patterns we could store with a network of this size.

**Figure 6:**
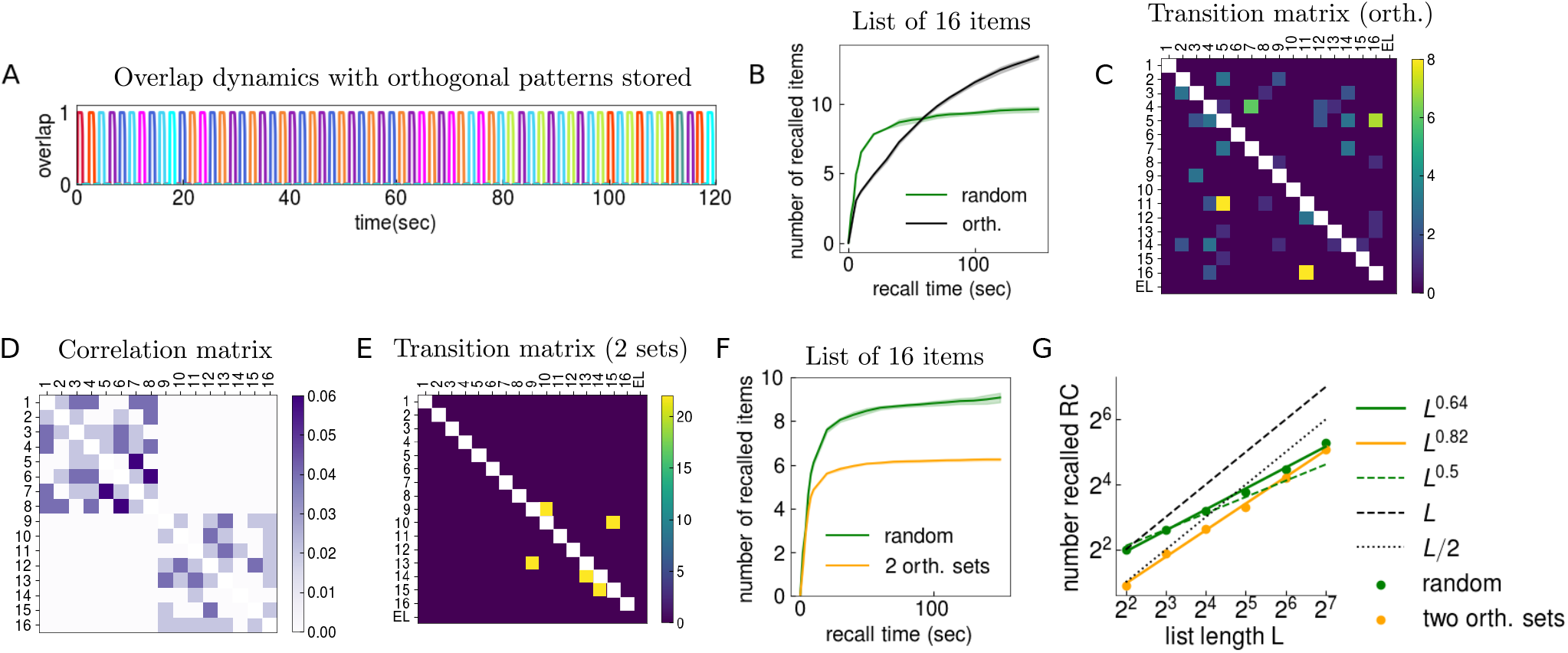
The structure of the stored memory set exerts a strong influence on the recall capacity. **(A)** We study our model with the storage of orthogonal memory patterns. Our network displays latching dynamics, hopping from memory to memory. This is shown through the overlap, or the correlation between the network state and a given memory, as a function of time (the different colours correspond to different memories). However, in this case, to be contrasted with the network storing random patterns, the network does not enter limit cycles, although there are bouts in the dynamics in which it recalls short sequences repeatedly. **(B)** The recall capacity as a function of time when orthogonal patterns are stored in the network (black curve). Compared to the random memory patterns (green curve), the network can recall all of the memory items in the list. **(C)** The transition matrix for one trial when orthogonal patterns are stored in the network. Compared to the random case (Fig. 5E), the network samples transitions more homogeneously. **(D)** We study our model with the storage of two sets of memories. Each set contains randomly correlated memories, but the two sets are orthogonal to one another. Pairs of memories belonging to the same set are randomly correlated but pairs in which memories belong to different sets are orthogonal to one another. The matrix expresses the correlation between the memories. The colourbar corresponds to the strength of the correlation. **(E)** The transition matrix for one sample recall trial. The network’s dynamics is entirely restricted to the second set (lower right block), as the network was cued with a memory item in this set. The colourbar corresponds to the number of transitions that have occurred. **(F)** The recall capacity is also limited in the case of the two-set model. In orange is the recall capacity as a function of time for the network having stored two sets of orthogonal patterns, and in green is the recall capacity for randomly correlated patterns, same as in Fig. 5D. **(G)** The recall capacity of the two-set model is bounded by the set size. Albeit with a different exponent, this recall capacity is still a power function of the list length (orange curve), to be compared with that of randomly correlated memories (green curve).

In order to test the behaviour of the model with more structured memory sets, we stored two orthogonal sets of long-term memories in the network. Each memory set contains randomly correlated memories, but the two memory sets are orthogonal to one another. Each list is comprised of memories from both memory sets, with half of the stimuli from one memory set and half from the second. Therefore pairs of memories belonging to the same memory set are randomly correlated but pairs of memories belonging to different sets are orthogonal to one another (Fig. 6D). In this case, inter-set transitions do not occur (Fig. 6E), and the dynamics is restricted to the set in which the network is initialised (Fig. 6 D, see also Fig. S6). Therefore the recall capacity is bound by the size of a single set (Fig. 6F). Finally, we find that the power law scaling holds also across the two-set model (Fig. 6G). Our model therefore predicts that although the statistics of the encoded memories has a dramatic effect on the recall capacity, the power law scaling, instead, may be expected to hold across more structured memory sets.

## 3 Discussion

In past decades, many models have studied the putative mechanisms behind the extensive information storage capacity of the human brain, and have advanced our understanding of the processes involved in memory storage [35–37, 9, 57–62]. However, it remains to be understood why recall of this information can be so limited, even when storage is guaranteed, as revealed by free recall experiments. We have shown how a simple network, with a biological plausible learning rule, can display recall properties such as those shown by human subjects. In our model, recall scales as a power law of the list length, similar to what has been found and recently corroborated for free recall of very long lists. This limited recall is attributed to the cycles the network enters into, hindering the recall of new items [31]. Our network may be thought of as implementing the abstract “algorithmic” model based on sparse random matrices, recently found to provide a parameter-free fit to human recall data [6]. In that class of models, transitions are made between items of highest similarity. Although we do find that transitions are also guided by correlations, they do not always follow the highest one, particularly when noise is present. Therefore, additional mechanisms, such as top-down attention may be modelled in order to stop recall after several items start to be repeatedly recalled. Another possibility may be the influence of slow inhibition [63], not taken into account in our model but that can be taken into account in more realistic network models comprising both excitatory and inhibitory neurons.

The storage/recall discrepancy in human memory recall may be partially resolved when taking into account the role of context, comprising different types of cues. For instance, even in a paradigm as simple as free recall, participants express various biases in their recall trajectories, biases that may reveal fundamental mechanisms meant to guide recall in natural settings outside of the paradigm. Many models have expressed context as an abstract component by itself, but it is not always clear what exactly they comprise [64]. We do not explicitly model context, but it may be thought to be provided by neuronal and synaptic processes that provide cues, for example correlations between representations provide semantic cues, whereas the temporal proximity of items allows for inter-item associations to take place, providing temporal cues that guide the recall trajectory. However, in order to achieve contiguity at multiple time scales, as has been found in experiments, and in particular across trials in multi-trial free recall, some form of temporal contextual cues may be necessary [65].

In our model, the addition of uncorrelated noise to the activity of neurons allows the network to recall more items by freeing the dynamics to explore beyond the limit cycle, a finding that may seem counterintuitive, given the canonical idea of the role of noise in neural networks. However, it must be stressed that the amounts of noise considered here are very small (*ε* = 0.05, not more than 5% relative to the signal the neurons receive from the recurrent connections). In support of this idea, it has been found that a targeted pulse of transcranial magnetic stimulation can lead to the reactivation of an item in human subjects, after the active representation of a presented item in memory drops to baseline [66].

We make no distinction between short-term and long-term stores, and effects traditionally attributed to them, such as the primacy and the recency effect emerge from the plasticity rules operating at different timescales, in the same network. The primacy effect is attributed to the spontaneous reactivations during interstimulus intervals, similar to what has been proposed before [67]. Such reactivations are not to be interpreted as rehearsals themselves, but may be read out by systems that express rehearsal, such as the phonological loop [68]. In support of this idea, in a free recall experiment with single cell recording in humans, verbal reports of memories of specific episodes at the time of free recall are preceded by selective reactivation of the same hippocampal and entorhinal cortex neurons [69]. Moreover, endogeneous reactivations have been found to occur in human subjects when engaged in unrelated activities through fMRI, and the extent to which they occur has been found to be correlated to later memory performance [70], in line with our model expressing primacy.

Finally, our results may provide predictions for future experiments. We predict that a small amount of noise, e.g. weak random stimuli or distractors should increase recall. Moreover we predict that the memory set structure plays an important role in guiding the transitions, and its manipulation has dramatic consequences on the recall capacity. Most studies of free recall have been performed with randomly correlated lists, and our results suggest the imperative to study more structured stimuli sets.

An interesting extension of this work would consider the case of multi-trial free recall, concerned with the the question of learning in a more general sense. For instance it has been found that when the order of word presentation is kept constant across multiple lists, the contiguity effect becomes progressively stronger. Conversely, when the order of word presentation within each list during list learning is randomised on each trial, the temporal contiguity effect progressively decreases, whereas the semantic effect increases across learning trials [71, 72]. Our study is beyond the scope of multi-trial free recall, but the mechanisms at play in our model can be reasonably expected to be consistent with this set of findings. In this spirit, a most intriguing puzzle that remains to be solved is how the rich structure of semantic associations in human memory arises partially as a result of the repeated exposure to items in temporal proximity [73–76].

In this contribution, we presented an attractor network model that recapitulates several qualitative and quantitative aspects of free-recall. Notably, only features such as a plausible learning dynamics, adaptation of neural activity, and global inhibition are necessary for the emergent dynamics of the network. Our model highlights the particular relevance of temporal and semantic correlations in guiding the recall trajectory, and stresses the importance of the memory set structure as a determinant factor of the recall capacity.

## 4 Methods

### 4.1 Simplified Model

We consider an attractor network of *N* binary neurons in which *p* sparse memories are stored. The memories are *N* bit vectors denoted by 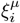, where *μ* = 1…*p* and *i* = 1…*N*. The 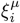 are randomly distributed and take the values 0 and 1 with the constraint that 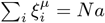 for any *μ*, where *a* is the fraction of active neurons in any given encoded patterns (the sparsity of the patterns). The long-term memories are encoded in the synaptic weights *J_ij_* according to the covariance rule

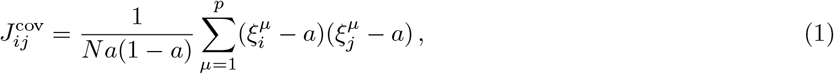

Up to now, this model is basically the one studied in [45]. We now assume that the neurons belonging to a set of *L* random patterns, out of the total of *p*, receive an additional fixed component *δJ* to their ingoing and outgoing weights, such that

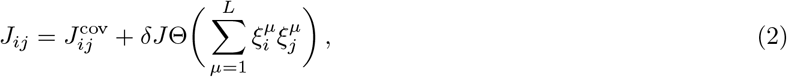

where Θ(.) is the Heaviside function. The update in a time step *dt* of the state of the neurons is performed asynchronously. At every step, all neurons are updated in random order. For neuron *i*, first we compute its local field as

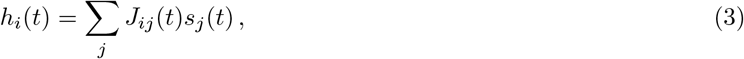

where *s_i_*(*t*) is the activity of neuron *i*. The local field is integrated by a rate variable *r_i_*:

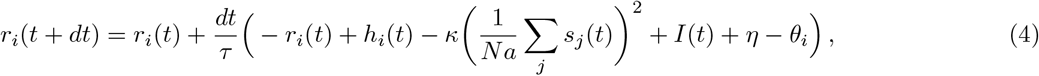

where, in addition to the depression provided by the covariance learning rule (ensuring a higher storage capacity [45]) there is an additional inhibitory component which is a quadratic function of the mean activity of the network [77], and proportional to the parameter *κ*. *I*(*t*) is the external input used to cue the network a list item, and *η* is a small random noise uniformly distributed within [−*ε, ε*]. The adaptive threshold *θ_i_*(*t*) represents neuronal fatigue, and is updated following

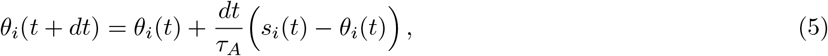

where *τ_A_* is the adaptation timescale. Finally, the activity is updated as

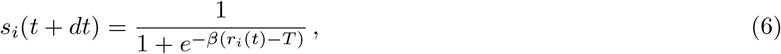

where *β* is the neuronal gain and *T* a fixed neuronal threshold. The integration time step is chosen to be *dt* = 0.01 seconds. Recall is measured through the overlap, defined as the correlation between the network state and each of the stored patterns

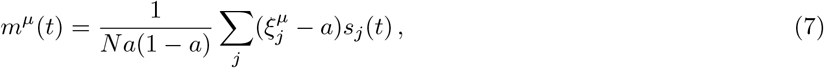

normalised so that it is 1 when the network state and a given memory are aligned. The neural variables *h_i_*, *r_i_*, *s_i_*, and *θ_i_* are all initially set to zero. The parameters used throughout the simulations can be seen in Table 2.

**Table 1:**
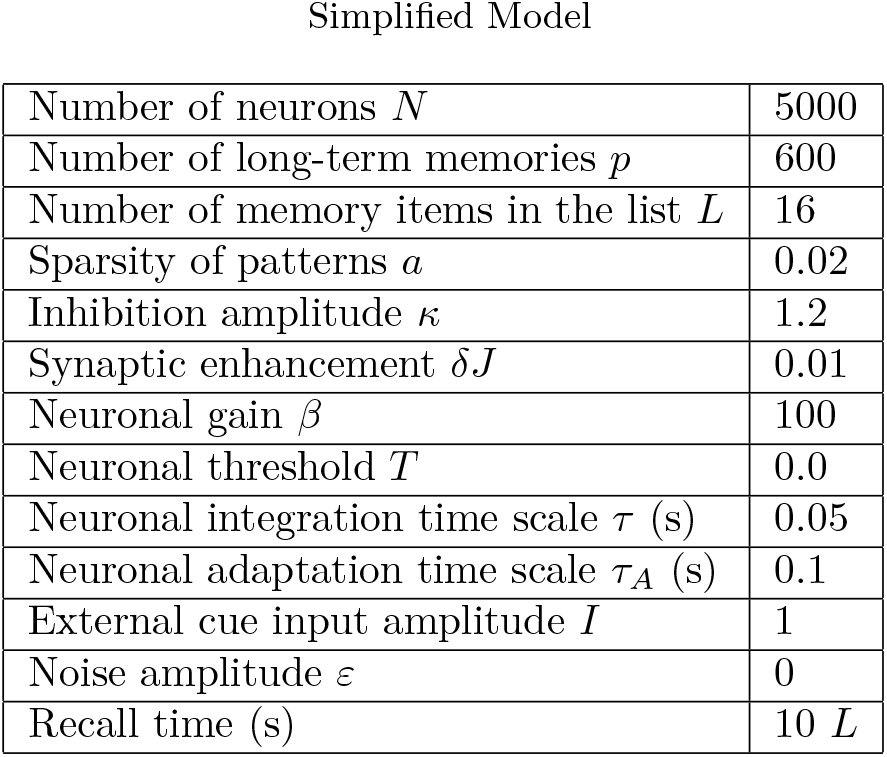
Simulation parameters of the simplified model, when not explicitly mentioned.

|  |  |
| --- | --- |
| Number of neurons $N$ | 5000 |
| Number of long-term memories $p$ | 600 |
| Number of memory items in the list $L$ | 16 |
| Sparsity of patterns $a$ | 0.02 |
| Inhibition amplitude $\kappa$ | 1.2 |
| Synaptic enhancement $\delta J$ | 0.01 |
| Neuronal gain $\beta$ | 100 |
| Neuronal threshold $T$ | 0.0 |
| Neuronal integration time scale $\tau$ (s) | 0.05 |
| Neuronal adaptation time scale $\tau_A$ (s) | 0.1 |
| External cue input amplitude $I$ | 1 |
| Noise amplitude $\varepsilon$ | 0 |
| Recall time (s) | 10 $L$ |

**Table 2:** Simulation parameters of the full model, when not explicitly mentioned.

|  |  |
| --- | --- |
| Number of neurons $N$ | 1000 |
| Number of long-term memories $p$ | 60 |
| Number of memory items in the list $L$ | 10 |
| Sparsity of patterns $a$ | 0.05 |
| Baseline facilitation $U$ | 0.5 |
| Inhibition amplitude $\kappa$ | 0.25 |
| Autoassociative learning rate $\alpha$ | 0.032 |
| Forward heteroassociative learning rate $\alpha_f$ | 0.01 |
| Backward heteroassociative learning rate $\alpha_b$ | 0.005 |
| Neuronal gain $\beta$ | 100 |
| Neuronal threshold $T$ | 0.03 |
| Neuronal integration time scale $\tau$ (s) | 0.05 |
| Neuronal adaptation time scale $\tau_A$ (s) | 0.1 |
| Synaptic facilitation time scale $\tau_F$ (s) | 7.0 |
| Neuronal activity trace time scale $\tau_S$ (s) | 5.0 |
| Stimulus presentation time $T_S$ (s) | 2. |
| Interstimulus interval $T_{ISI}$ (s) | 1. |
| External cue input amplitude $I_{cue}$ | 1. |
| Noise amplitude $\varepsilon$ (distractor) | 1. |
| Recall time (s) | 10 $L$ |

### 4.2 Full Model

In the full model, learning is made explicitly dynamic. We assume that the synaptic weights *J_ij_* can be expressed as

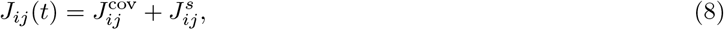

where the potentiation 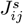 in addition to the Hebbian weights *J*^cov^ is driven by three different contributions. First, the instantaneous co-activity of the pre- and post-synaptic neurons, represented by the product *s_i_ s_j_*. This contribution alone would provide the dynamic Hebbian learning rule. Second, the activation of the post-synaptic neuron with a delay Δ*t* after the activation of the pre-synaptic neuron, that is *s_j_*(*t* – Δ*t*) *s_i_*(*t*); this is what we refer to as the *forward* heteroassociative learning. Finally, the activation of the pre-synaptic neuron after the post-synaptic, by a delay Δ*t, s_j_*(*t*) *s_i_*(*t* – Δ*t*); we termed this the *backward* heteroassociative learning. For these second two contributions, assuming that the delay is exponentially distributed, the average contribution would be

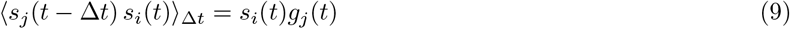

where *g_j_* is the *activity trace* of neuron *j*, defined as the exponential average of the past activity on a time scale *τ_S_*:

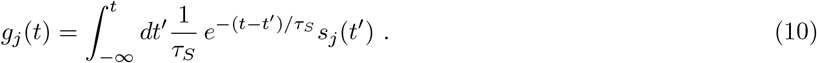

Analogously for *i* and *j* interchanged. This quantity can be seen to satisfy

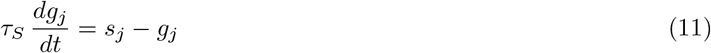

We note in passing that if *τ_S_* = *τ_A_*, as in [78], this equation is equivalent to the one for the adaptation variables *θ*. If the activity trace coincided with the adaptation variable, it would be appealing because 1) the complexity of the computation in the biological network would be reduced and 2) it may provide an additional physiological role to the neuronal fatigue, as the stress that strengthens the synapses.

We also introduce a regularisation term that prevents the synaptic weights from becoming infinite upon sustained co-activation of pre- and post-synaptic neurons. We add a linear “degradation” term, i.e. a negative contribution proportional to the weight, akin to Oja’s rule, which is designed to conserve the norm of the weights vector.

Taken together, all these contributions, implemented asynchronously neuron by neuron, yield, for the ingoing weights 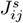

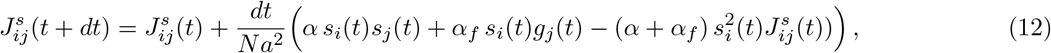

while for its outgoing weights –that is the 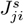 where for *j* post-synaptic to *i*– we have

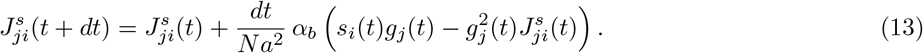

During the whole network update, neurons’ states are updated randomly without replacement, therefore each neuron is updated only once. The corresponding ODE writes

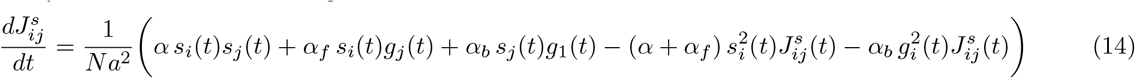

In addition, we also include a presynaptic facilitation *u* of the weights [79, 80]

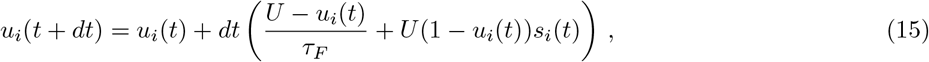

where *τ_F_* is the facilitation timescale and *U* is the baseline facilitation.

The input to a neuron is given by

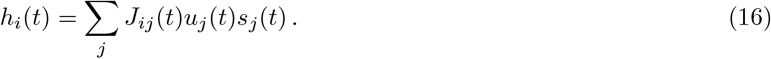

The field *h_i_* calculated in Eq.(16) is then used to update the activity of neuron *i* according to the rules in Eqs. (3–6). This input is integrated by a rate variable *r_i_*:

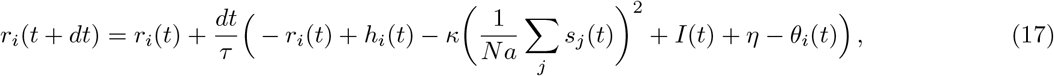

where *I*(*t*) is the input used to cue the network with list items. Initially, the neural variables *h_i_, r_i_*, *s_i_, θ_i_* are all set to zero. *u_i_* is set to its baseline value of 0.5, while the initial value of *g_i_* is set to 0.05 for all neurons. We sequentially present memory items to the network for a duration of *T_S_* followed by interstimulus intervals of *T_ISI_*. After list presentation, we assume that learning has taken place, and there are no more weight changes. We then let the network recall the list spontaneously for a time equal to *T_R_* = 10*L* seconds. The parameters used throughout the simulations can be seen in Table 2.

### 4.3 Conditional Response Probability

The algorithm used to compute the lag-CRP, designed by Kahana is explained in [48]. We use the codes provided in http://memory.psych.upenn.edu/CRP. In order to compute one CRP curve, we run 480 trials in which the set of *p* patterns is kept constant, and where the set of *L* patterns is randomly selected (without replacement) from the *p*. Then we repeat the same procedure 5 times, with different sets of *p* patterns. The errorbars in all curves correspond to the standard deviation of the 5 curves.

In order to compute the semantic-CRP, we repeat exactly the same procedure, but instead of the lag, we consider the correlation between patterns *C^μν^*, defined between pattern *μ* and *ν* as

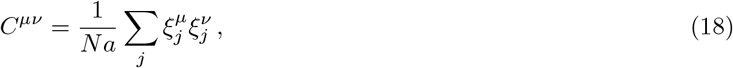

that takes discrete values due to the fact that *N* is finite.

## 5 Supplementary material

### 5.1 Mean field analysis of the simplified model

In this section, in order to better understand the dynamics of the spontaneous reactivations, and the conditions under which they arise, we develop a mean-field theory of our model under several simplifying assumptions. The first assumption that we make is that we neglect *J^s^* in Eq. (2). The addition of *J^s^* breaks the symmetry among the *p* patterns, by favouring the activation of the *L* patterns coding for items in the list. This restricts the dynamics of the network only to the items in the list – save for the few exceptions discussed in the main text (the excursions). However, this term does not entail a qualitative difference in the several dynamical regimes, brought about mainly by firing rate adaptation and global inhibition. For this reason, and in order to simplify the analysis (meant to understand qualitative features of the dynamics), in this section we consider synaptic weights given the first term of Eq. (2) only, that is the Hebbian weights *J*^cov^.

We start by noting that, in continuous time, Eq. (6) at steady state can be written as

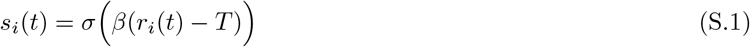

where *r* is the activity rate, *T* the static neuronal threshold, *β* the neuronal gain, and where the sigmoid function *σ* is defined as

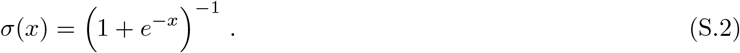

The evolution of the activity rate *r* is given by

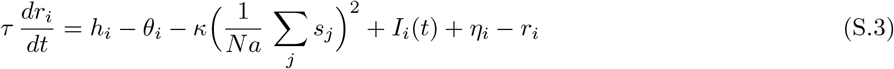

where *τ* is the neuronal time scale, *h* is the total input to a neuron, *θ* is a variable threshold modelling firing rate adaptation, *κ* is the strength of the global inhibition, *I* is a uniform external input, and *η* is a uniformly distributed random input. The evolution of the adaptive threshold is given by

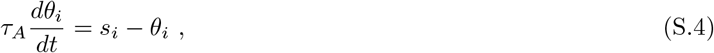

where *τ_A_* is the adaptation time scale. The input field *h_i_* is given by

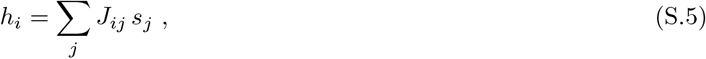

that is, the weighed sum of presynaptic activities, with prescription weights

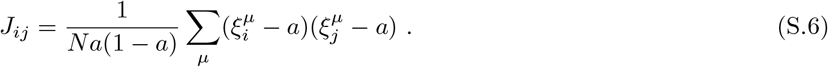

The *ξ*^*μ*^s correspond to *N*-bit vectors representing the memories with sparsity (*a* i.e. the probability that neuron is active in a given pattern). Equations (S.3) and (S.4) are the continuous, differential form of Eqs. (17) and (5), respectively.

If we assume that *τ* ≪ *τ_A_*, over time scales which are comparable to or larger than *τ_A_*, the variables *s* and *r* are adiabatically slaved to the dynamic thresholds *θ*: as *θ* changes in time, *s* and *r* quickly relax to the quasi-steady state specified by the current *θ*. In this approximation, as the intermediate variable *r* does not directly influence *θ*, but does so only by defining the quasi-steady state of the activity *s*, it can be removed from the analysis, therefore obtaining the following set of equations

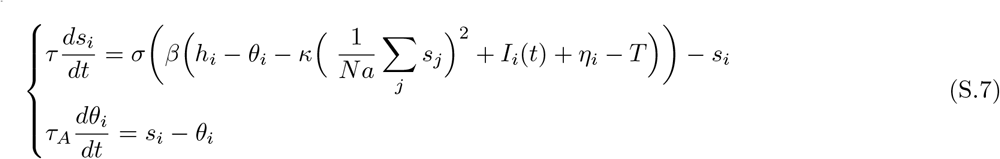

where *τ* ≪ *τ_A_* sets the time scale of relaxation of *s* to the steady state, and reflects the underlying activity rate.

In order to describe the transition dynamics of the network from pattern to pattern, and since we are interested in solving the set of coupled equations given by Eq.(S.7), we need to derive the equations for the activity vectors *s* and the corresponding thresholds *θ* as projected onto each individual patterns: for the pattern *μ* we define the corresponding *activity overlap* and *adaptive threshold overlap* as

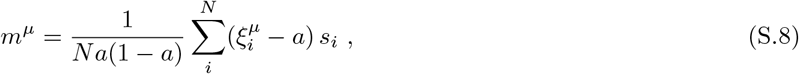

and

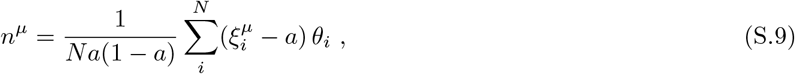

respectively. We therefore apply this projection to both equations in Eq(S.7), and derive approximate differential equations for the overlap variables. As the projection is linear, the left-hand side of Eq. (S.7) trivially yields the time-derivatives of the overlap variables. The right-hand side of the equation for the adaptive threshold is linear, so *s* and *θ* are simply replaced by their respective projections, the overlaps *m* and *n*. The non-trivial part of the calculation lies in the equation for the activity that involves the non-linear activation function *σ*, and for which approximations are made.

First, we notice that the input field of neuron *i* in Eq. (S.5) can be written as

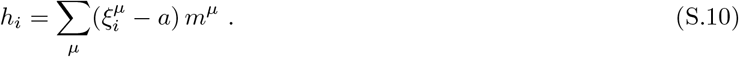

Then, we also notice that when the sparsity *a* is small (i.e. a small fraction of neurons are prescribed to be active in each pattern, and more precisely when *a p* is smaller than or comparable to *N*) to a first approximation, we can assume that there is negligible overlap between any two patterns. Therefore, sums over neurons can be split into sums over the neurons that belong to each individual pattern. In particular, the total activity that enters the global inhibition term in Eq. (S.7) can be expressed as

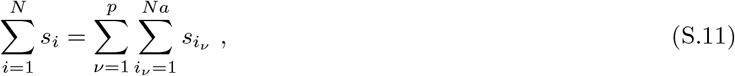

where *v* indicates the pattern, and *i_ν_* the subset (partition) of neurons that are active in pattern *v*. Sums of neuron-dependent quantities over the neurons of a specific pattern only, in turn, can be approximately written as the overlap of that quantity with the pattern of interest. Specifically, for the activity or the adaptive threshold variables, we have

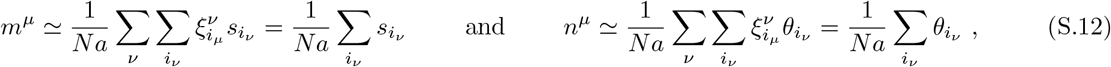

which follow from Eqs. (S.8) and (S.9) in the approximation where *a* ≪ 1 and for non-overlapping patterns, for which 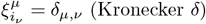. It follows that the global inhibition term is expressed in terms of the overlaps as

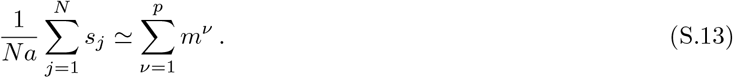

Momentarily neglecting the fixed threshold *T*, the noise *η_i_* and the external input *I*, these observations allow us to write the equation for the activity overlaps *m* in an approximate form as

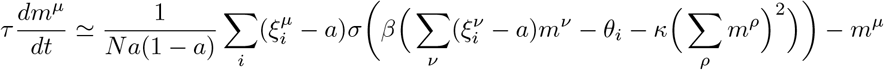

Again, in the approximation where *a* ≪ 1, the sum is restricted on the set of neurons which are active in the pattern of interest only, so

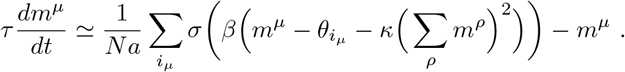

The only remaining neuron-dependency is carried by the adaptive threshold *θ_i_*. The right-hand side can be seen as the average over the distribution of *θ* for neurons in pattern *i*. If the distribution *θ* is concentrated enough, we can assume the distribution of the *σ* is concentrated near 0 or 1. So, despite the non-linearity of the sigmoid function, we can “take the average” inside *σ*, and by using Eq. (S.12) we can write

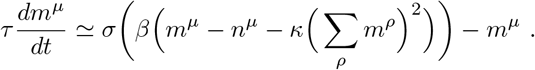

The error in this last approximation, when the distribution of *θ* overlaps with the range of values where *σ* varies substantially, can be neglected for a two reasons. First, the overlap can be explained by a smaller value of *β* –now a phenomenological parameter– which increases the width of the “switch” interval. Second, when this happens, it does only for a short amount of time, as the thresholds would drive the activity overlaps to perform a rapid switch, and therefore quickly moving the problematic interval away.

There are some terms that have been neglected and that are worth discussing. First, the correlation between patterns –that is the neurons co-active in more than one pattern– may not be negligible. In fact, when patterns represent items, these correlations may express the semantic relation between them, and in the main text we show that such correlations are important in guiding the transitions in recall. Second, while we isolated relevant contributions to the terms that appear as arguments of the activation function *σ* in Eq. (S.7), we may also include the neglected terms by the addition of a random term *η^μ^* with a constant mean *γ* and fluctuations uniformly distributed within [−*ε, ε*]. This phenomenological noise term can be re-defined in order to absorb the noise *η_i_*, which had been neglected above. Similarly, the fixed threshold *T* can be absorbed into the phenomenological constant *γ*.

In conclusion, we can write a mean-field approximation of Eq. (S.7) as

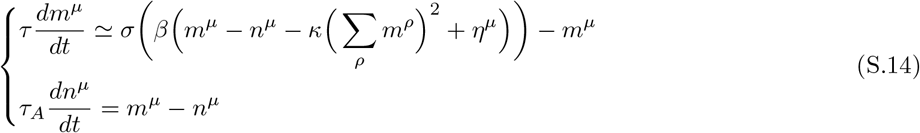

The external input *I_i_* with arbitrary neuron-dependence cannot be included in this approximation. However, if one wants to model a constant input given only to neurons that code for a given pattern, the same arguments hold as for the treatment of the adaptive threshold *θ_i_*, where *I_i_* would be replaced by its average over the relevant neurons. This can be included by an addition of an extra contribution to *γ* only for the pattern of interest.

It is important to note that, even when the correlation between patterns is negligible –which is one of the assumptions in the analysis above– the presence of the global inhibition term, weighed by the parameter *κ*, provides interdependence, hence “interaction” between the overlaps with different patterns.

#### 5.1.1 Phase diagram for the single-pattern case

When the strength of the inhibition *κ* is vanishing, Eq. (S.14) prescribes an independent dynamics for the individual overlaps. Therefore, we focus here on the dynamics of the overlap *m* ≡ *m*^1^ of a single pattern *ξ* ≡ *ξ*^1^.

As the time scale for the state dynamics *s* is much shorter than that of the adaptive threshold *θ*, when the noise is negligible, *ε* = 0, we can solve for the quasi-steady state of *m*, i.e. its stationary value given a “frozen” *n*

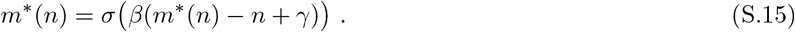

When a small noise term is introduced, this is achieved on average.

If *β* ≫ 1, the sigmoid function *σ* is approximated by the Heaviside function Θ, defined as Θ(*x* ≥ 0) = 1 and Θ(*x* < 0) = 0, so that the solution *m\** can only be 0 or 1. The steady state solution depends on the value of *γ, n* and the initial condition of *m*. If *n* – γ < 1, then both 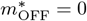 and 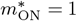 are stable solutions, and they are attained depending on whether the initial value of *m* is below or above *n* – *γ*, where the unstable solution *m*\* = 1/2 is found. Instead, if *n* – *γ* > 1, only 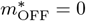 is a solution. Large but finite values of *β* change these steady states slightly below 1 and above 0, respectively.

One can now consider the dynamics of *n* in the adiabatic approximation wherein *m* is replaced by its steady-state value *m*\*(*n*).

##### Inactivation

Suppose that the overlap is initially high enough, so that *n* – *γ* < *m*; then, *m* quickly attains the value 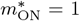, then followed by *n* on a slower time scale. However, if *γ* < 0, when *n* reaches the critical value 1 + *γ*, the overlap immediately switches to the stable fixed point of Eq. (S.15), 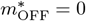, which becomes the new (instantaneous) attractor of the dynamics for the adaptive threshold. Instead, if *γ* > 0, the state *m* = 1 is always a stable attractor, and the adaptive threshold reaches asymptotically the steady state *n* = 1.

Finite temperature values, that is *β* < ∞, contribute to reducing the critical value of *n* which causes the “OFF switch” of the overlap. In the condition of coalescence between the stable and unstable solutions of Eq. (S.15), at first order in *β*^−1^, we have 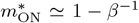 –found by solving the degeneracy condition where derivatives with respect to *m* on either side of the equation match, giving *m* – *n* + *γ* – *β*^−1^ log *β*, which is then used to solve Eq. (S.15) approximately. The critical value of *γ* for which an OFF switch can occur is found by substituting into the degeneracy condition the fixed-point value 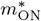 and the maximal value of *n*, that is again 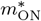. This trivially gives us the upper critical value of *γ*, *γ*^+^, as a function of the inverse temperature *β*,

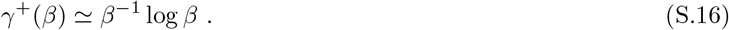

For *γ* < *γ*^+^, the value of the adaptive threshold that triggers the OFF switch is given by

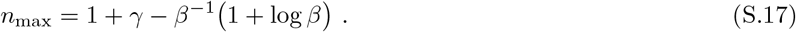

For *γ* > *γ*^+^, instead, *n* reaches the the steady state 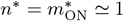.

##### Activation

When *m* < *n* – *γ*, either as initial condition of the system or upon an inactivation event, Eq. (S.15) prescribes that the overlap reaches fast the quasi-steady state 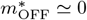. It follows from Eq. (S.14) that *n* will also approach vanishingly small values. As it does, the OFF state may become unstable and the overlap may perform an “ON switch”. Similarly for case of an inactivation, this instability, may or may not be reached depending on the values of *β* and *γ*. An analogous calculation as the one for the inactivation events shows that, for a large but finite *β*, it is it is possible to find a lower threshold *γ*^−^ such that for *γ* > *γ*^−^ an ON switch can occur. This is found to be

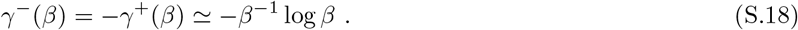

The value of the adaptive threshold triggering the ON switch is also derived to give

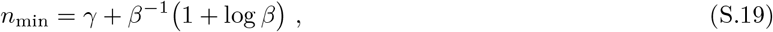

for *γ* > *γ*^−^. Instead, for *γ* < *γ*^−^, no activation can occur, and *n* reaches the steady state 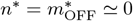.

##### Oscillations

As we have calculated the turning points for the dynamics in *n*, we can estimate the time intervals during which the overlap is continuously ON after an activation event, *t*_ON_, or OFF after an inactivation event, *t*_OFF_. By analytically integrating the equation for *n* with the corresponding value of 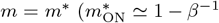 for *t*_ON_, 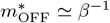 for *t*_OFF_), the ON/OFF times can be calculated analytically as the times between the two turning points *n*_min/max_,^1^

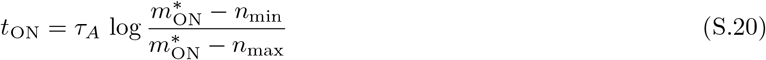

and

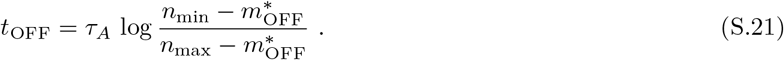

We note that, although the equation for *n* is linear, when we solve it in the adiabatic approximation for *τ* → 0, it becomes non-linear, as *m* should be replaced by its instantaneous steady state *m*\*(*n*). Still, the linear approximation is good, because *m\** is approximately constant for a high *β*. In an initial transient, when the switch has just occurred, *m*\* is either 0 or 1, which are the initial “target” values for the dynamics of *n*. As *n* approaches the critical value for the switch, however, we showed that the target has a correction of order *β*^−1^, that is 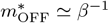 and 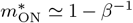, where the dynamics sees a slow-down and therefore gives the larger contribution to the inter-switch times. Therefore, we use the corrected values as the target for the dynamics of *n*.

One can check, for consistency, that *t*_ON_ and *t*_OFF_ are mathematically defined, for a given *β*, when *γ* > *γ^−^*(*β*) and *γ* < *γ*^+^(*β*) respectively. When both are defined via Eqs. (S.20) and (S.21), an oscillatory behaviour is observed, with *n*_min/max_ as lower/upper turning points. Where *t*_ON_ (*t*_OFF_) is not defined, the overlap is always approximately unity (vanishing), and we refer to this as a ferromagnetic (paramagnetic) behaviour, in analogy with spin systems. The time average of the overlap then is simply

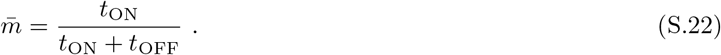

This analytic result is shown for the deterministic case (Fig. S2 A) and matches the simulation results for large values of *β* (Fig.S2B).

##### Effect of noise

When noise is added to the system (*ε* > 0), simulations show that the oscillatory phase is widened (Fig. S1 A second row). This can be intuitively understood because random values of the noise may have a switch performed before *n* reaches the critical value *n*_min/max_ that we estimated analytically for the deterministic case. As a result, where we would expect a paramagnetic behaviour for *ε* = 0, we may instead have a period of random short-lived activity, triggered by noise at irregular intervals (*RA*, Fig. S1 A and Fig. S1B panel f). Analogously, where the deterministic system would be ferromagnetic, with noise we may instead have random inactivation events at random time (*RI*, Fig. S1 A and Fig. S1 B panel d). When oscillations are expected even at the deterministic level, with noise we observe oscillations with higher frequency (*DO*, Fig. S1 B panel e). This is again due to the fact that activation/inactivations occur before *n* could reach the critical values.

##### Effect of the inhibition on a single pattern

We can also see the effect of the addition of a global inhibition term, which contributes to reducing the amplitude of the oscillations (Fig. S1 second and third columns). Due to the quadratic form of the inhibition, the phase diagram is substantially changed only where the system is close to being ferromagnetic for *κ* = 0.

#### 5.1.2 Multi-pattern case

##### Role of inhibition

The multi-pattern case is the context where we best see the role of global inhibition. When *κ* = 0, the equations for the overlaps of different patterns are completely independent, and the dynamics can be understood simply in terms of the single-pattern case. Instead, when *κ* > 0, the global inhibition creates a *competitive* interaction among patterns. The activity of the network cannot span all patterns at a time, but fewer and fewer as *κ* increases. This has interesting staggering effects on the dynamics. For *κ* = 0, if activity and adaptive threshold overlaps of all patterns are close at some time, they remain close at all times (Fig. S3 A). Instead, when *κ* > 0, a small difference in the activation function experienced by different patterns gets quickly amplified because of the inhibition: the pattern with the highest *m* dominates at the expense of the others, until the corresponding adaptive threshold contributes to switching it OFF and other patterns may be retrieved. For instance, in the two-pattern case (Fig. S3 B), we see that after an initial transient the activity overlaps become counter-phased, for arbitrarily close initial condition. Interestingly, simulations show that the system gets locked into some quasi-stationary cycles, where activity overlaps are repeated in the same order.

##### Disturbance by a large noise

The mean-field model also captures the behaviour when a large amount of noise is introduced into the system. When there is no noise *ε* = 0, the mean field equations express latching dynamics (see Fig. S3 C). Instead, when *ε* =1, this perturbation is such that all patterns tend to have a finite average activity. However, the inhibition term becomes dominant when the number of patterns is high, and forces the average activity to be low (Fig. S3 D). In the main text, we described this as a model implementation of a distraction manipulation, which we showed to heavily impair the recall of any pattern.

**Figure S1:**
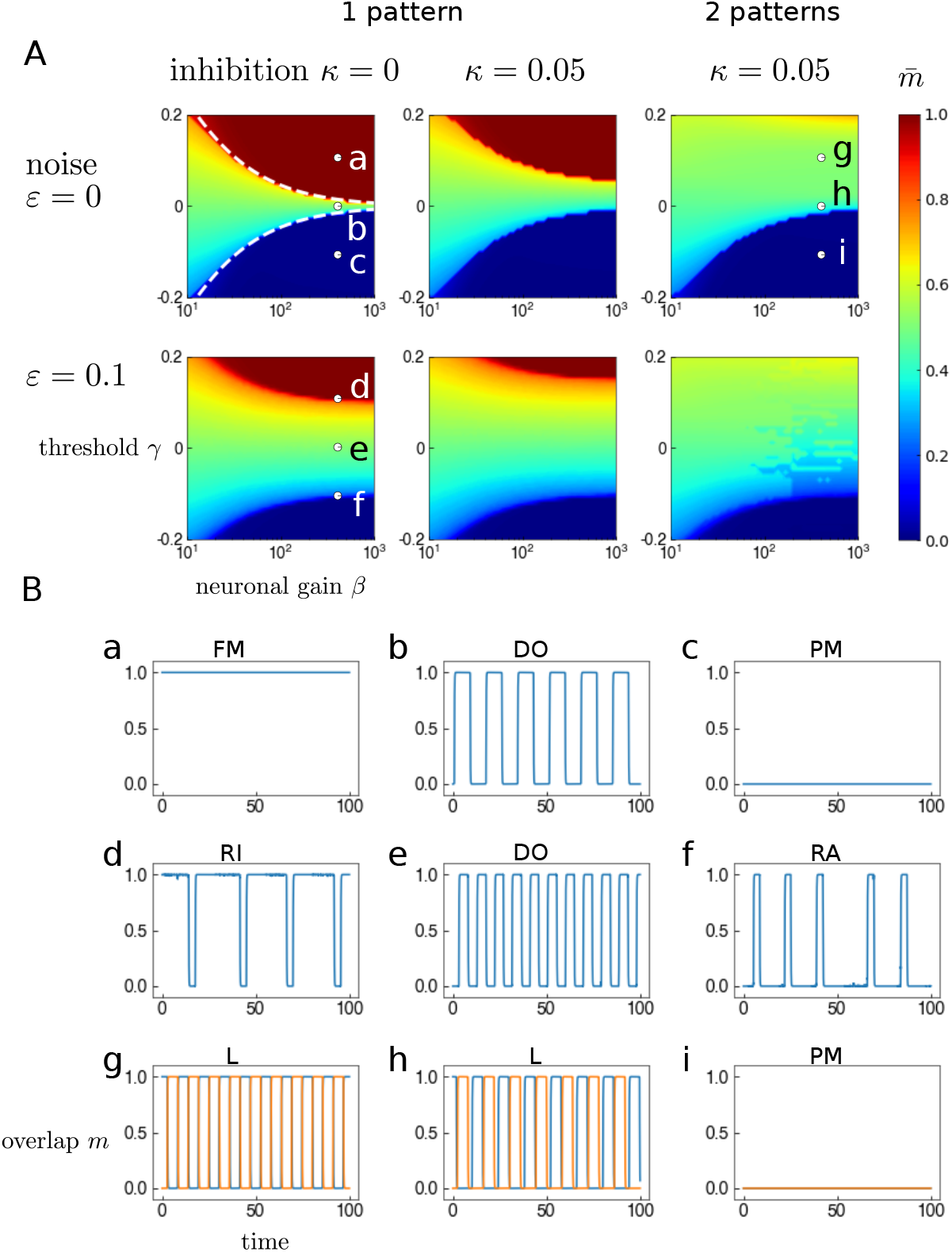
Different regimes of network operation. **(A)** Phase diagrams 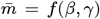 of the single-pattern and two-pattern case. The colourbar corresponds to the average over time 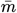 of the activity overlap, obtained through the numerical integration of the mean-field equations Eqs. (S.14). The *x* and *y* axes correspond to the neuronal gain *β* and threshold *γ*. Columns correspond to different values of the inhibition parameter, *κ* = 0 and *κ* = 0.05. For the deterministic case with no inhibition (*κ* = 0 and *ε* = 0), analytical estimates of the phase boundaries are plotted as white dashed lines – Eqs. (S.16) and (S.18)). Here, three phases are present: ferromagnetic (*FM*) 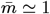 (dark red), paramagnetic (*PM*) 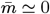 (dark blue), and oscillatory (*DO* – deterministic oscillations), marked by a, c and d, respectively. Sample trajectories of the overlap in these regimes (exemplified by these points) are shown in (B). Comparison between the left and middle panels shows the effect of global inhibition *κ*: due to its quadratic dependence on *m*, the paramagnetic phase 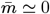 is substantially unchanged, while the ferromagnetic phase 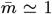 is gradually reduced upon increasing *κ*. With two patterns (rightmost column), the ferromagnetic phase disappears, at least in this range of the parameter *γ*. This can be traced to the competitive interaction between patterns (see text for more details). Rows correspond to two different values of the noise, *ε* = 0 and *ε* = 0.1. Noise widens the oscillatory phase. Noise also contributes to a similar effect by blurring the boundaries between phases (marked by d and f): random activations (inactivations) occur near the points where we would expect a transition between the oscillatory and the paramagnetic (ferromagnetic) phases. **(B)** Sample trajectories of the overlap, *m*, representative of all the qualitative behaviours observed in the phase diagrams in (A) and corresponding to the parameters marked a-i. The top row corresponds to the single pattern case without inhibition, *κ* = 0 and without noise *ε* = 0. The middle row corresponds to the case without inhibition and *ε* = 0.1. In this case, different behaviours can be observed, which show the signature of stochasticity in the system in regions of parameters otherwise characterised by *FM* or *PM*: random inactivation events when the system is typically ferromagnetic (*RI*), and vice versa, random activation events in a mainly paramagnetic regime (*RA*). The last row corresponds to the two-pattern case with *κ* = 0.05 and *ε* = 0. Blue and orange correspond to two different colours. The oscillatory phase in the multi-pattern case, where alternate activations between patterns occur, is labelled as latching (*L*).

**Figure S2:**
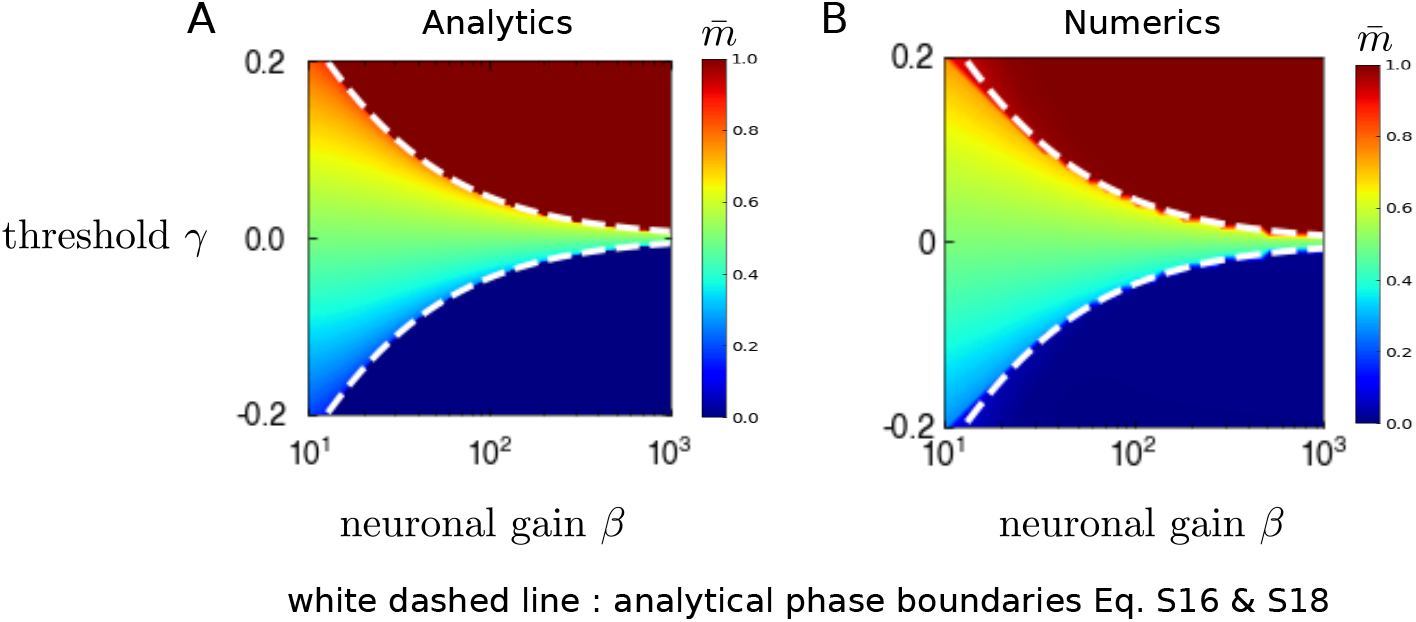
Comparison of the analytical result Eq. (S.22) for the noiseless case is in agreement with numerical integration of the mean-field equations. **(A)** Analytic phase diagrams for the deterministic case, Eq. (S.22). The colourbar corresponds to the overlap order parameter 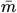. **(B)** Phase diagram obtained through the numerical integration of the mean-field equations, Eq. (S.14). The white dashed lines in both panels correspond to Eqs. (S.16) and (S.18). For large values of *β*, the analytical result Eq. (S.22) is in excellent agreement with numerical integration of the mean-field equations Eq. (S.14).

**Figure S3:**
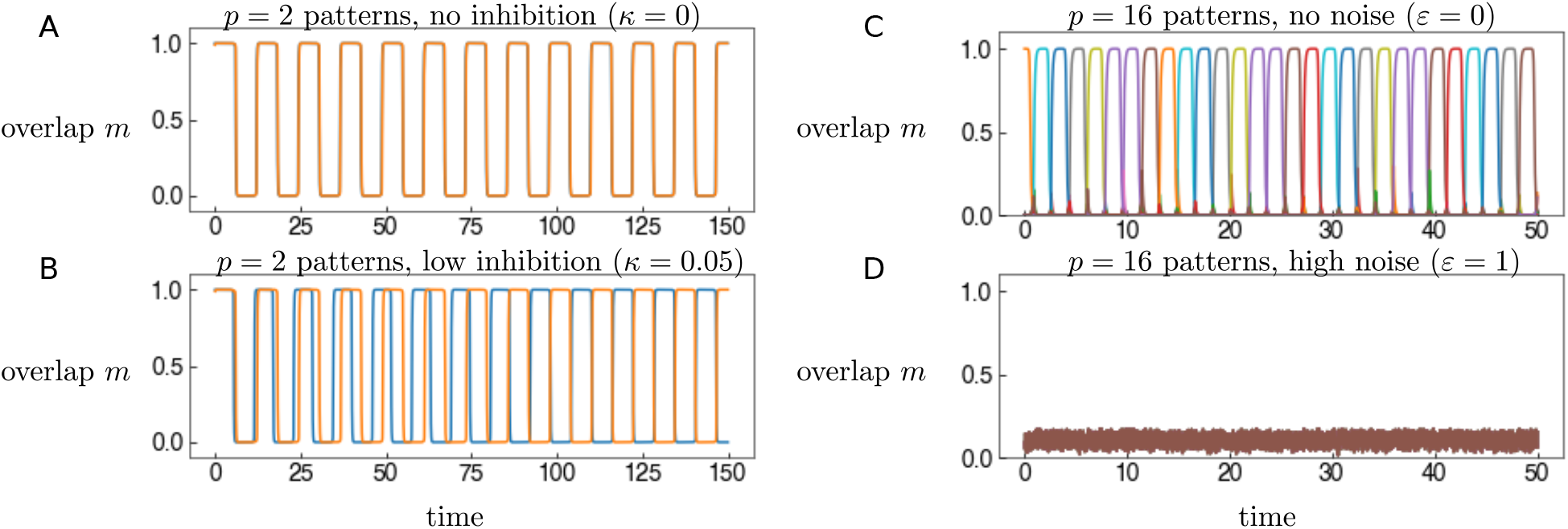
The roles of inhibition and noise. **(A)** Overlap as a function of time, obtained through the numerical integration of the mean-field equations Eq. (S.14) in the two-pattern case for *κ* = 0. The two patterns have very close initial conditions for both the activity and adaptive threshold overlaps (the latter not shown). Orange and blue correspond to the two patterns. The two overlaps are almost identical because patterns have identical dynamics, but a very small difference in initial conditions. **(B)** Same as in A), but with *κ* = 0.05. The addition of a small amount of inhibition triggers competition between the patterns increasing the initial unbalance. The trajectories of the two overlaps, while initially almost perfectly coincident, become staggered after a transient. The other parameters are *β* = 100, *γ* = 0 *ε* = 0. **(C)** Overlap as a function of time, obtained through the numerical integration of the mean-field equations Eq. (S.14) with *p* = 16. Different colours correspond to different patterns. Here *ε* = 0, i.e. no noise present. **(D)** Same as in A), but with *ε* =1. The presence of such large noise shuts down all activity, and sets the network in the paramagnetic regime.

### 5.2 Effect of learning rates and inhibition on the network dynamics

**Figure S4:**
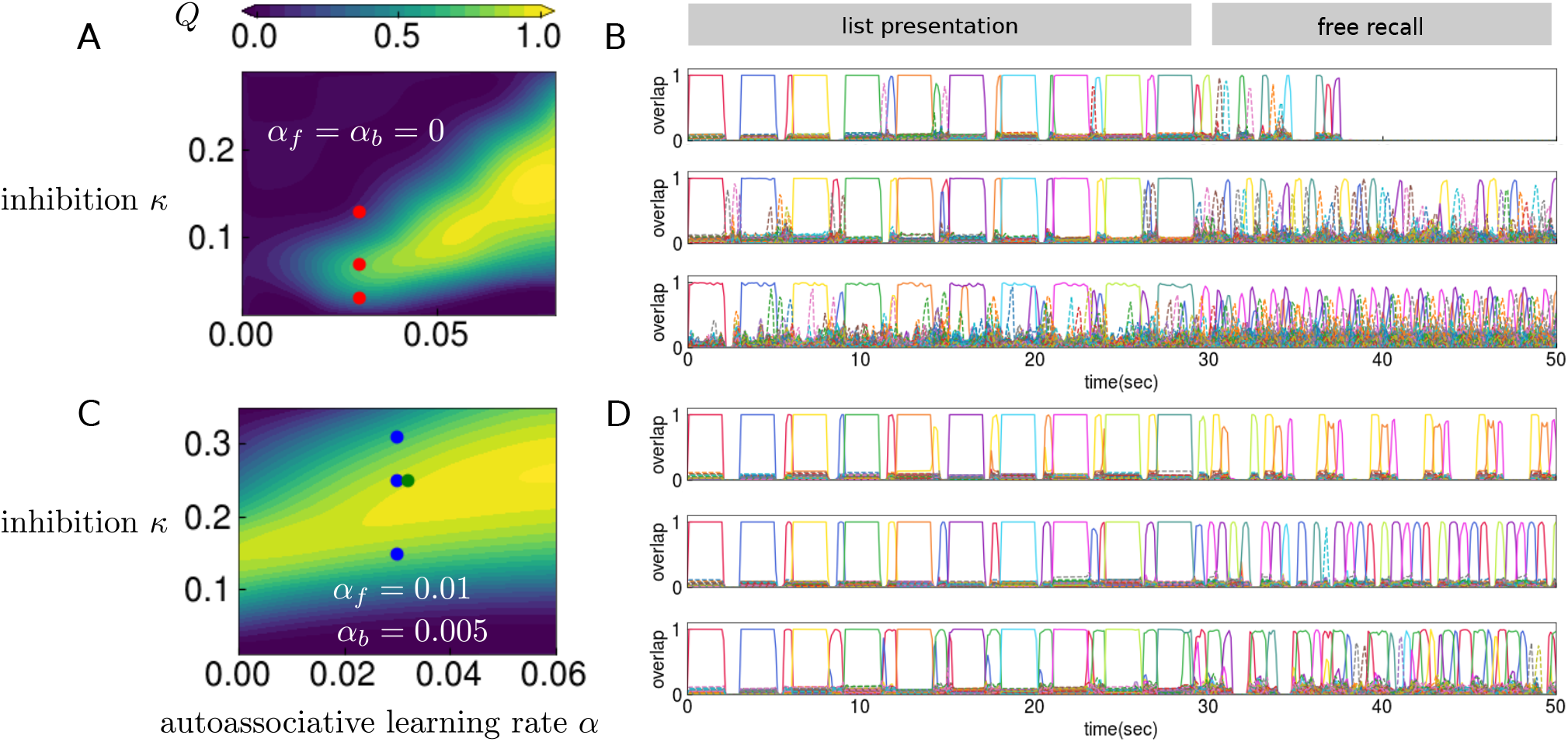
Sufficiently high learning rates and inhibition are necessary to simultaneously push the full model to reactivate continously as well as restrict its dynamics to recalling the list. We consider a model of long-term memory [45] equipped with dynamic auto- and heteroassociative weights, in addition to the baseline weights (details in Methods). *α* is the autoassociative learning rate, whereas *α_f_* and *α_b_* are the forward and backward heteroassociative learning rates. We simulate our model with a free-recall task: the presentation of an item corresponds to the addition of an external input field to all the neurons coding for that item, after which the network is left to freely recall items. Recall is measured through the overlap, or the correlation between the network state and a given memory, as a function of time (the different colours correspond to different memories). **(A)** Plot of a measure 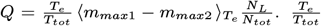 expresses the duration that the reactivations are sustained endogenously, relative to the duration of the whole simulation. 〈*m*_*max*1_ – *m*_*max*2_〉*_T_tot__* is the difference between the largest overlap and the second largest overlap, averaged over the entire sequence duration *T_e_*, expressing the discriminability of the recall sequences [41]. Finally, 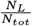 is the fraction of intralist items, relative to the total number of items recalled, expressing how well the network restricts its dynamics to the pool of *L* items in the list. The colourbar expresses this measure when both heteroassociative learning rates *α_f_ = α_b_* = 0. **(B)** The three individual trajectories of the overlap *m* correspond to the three values of the inhibition parameter *κ* marked in red in the phase diagram in (A). When inhibition is too low, the sequences are not well discriminable, and oftentimes multiple memories are recalled simultaneously (see also mean-field analysis and Fig. S3). When it is increased, sequences become more discriminable, but extralist items (shown in dashed coloured lines) still plague the dynamics. Further increasing the inhibition suppresses even intra-list memories (shown in uninterrupted coloured lines). **(C)** The same measure as (A) is plotted here, but for the forward heteroassociative learning rate *α_f_* = 0.01 and the backward heteroassociative learning rate *α_b_* = 0.005. In contrast to the previous case, a sufficiently high learning rate *α*, combined with sufficient inhibition *κ*, assures that sequences are discriminable (see also mean-field analysis and Fig. S3) and dynamics is largely restricted to the list, as shown in (D). The point marked in green is the value of the learning rate that has been used in all of the simulations of the *full model* in the main text. **(D)** The three trajectories of the overlap *m* in this panel correspond to the three values of the inhibition parameter marked in blue in the phase diagram.

**Figure S5:**
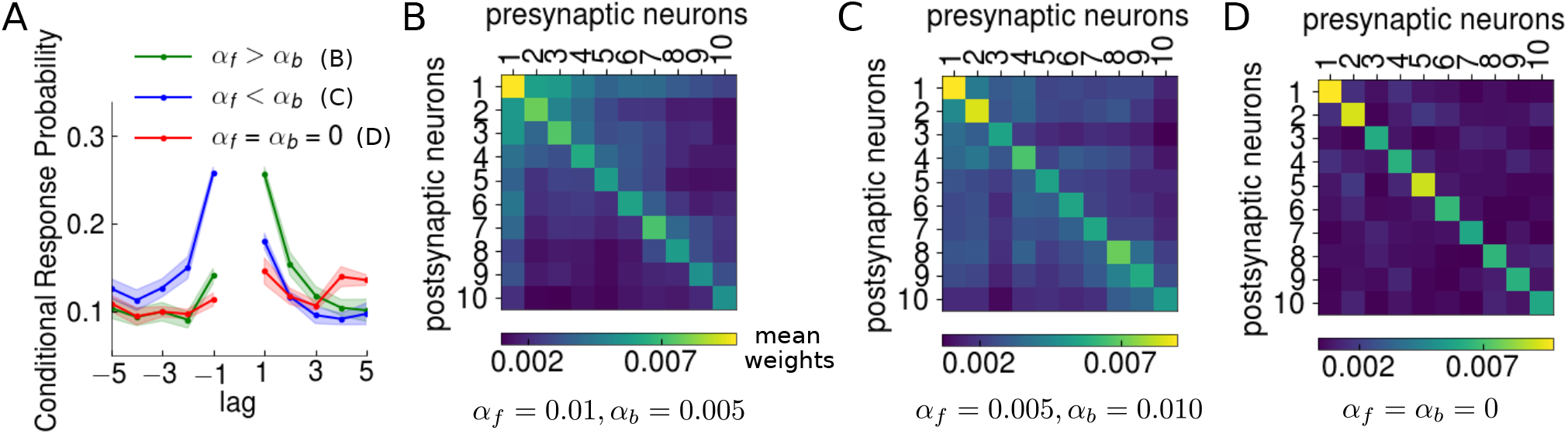
Conditional Response Probabilities with differing forward and backward rates in the full model. **(A)** The Conditional Response Probability is computed as explained in Methods. The contiguity and forward asymmetry of transitions is determined through different values of the learning rates. When learning is purely autoassociative, i.e. *α_f_* = *α_b_* = 0, there is no contiguity effect, and the asymmetry is minimal (red curve). When the forward heteroassociative learning rate *α_f_* is greater than the backward rate *α_b_* (green curve also shown in main text in Figs. 3 A, 3 C and 3E) both contiguity and forward asymmetry are present. Conversely, when this pattern is reversed, the asymmetry becomes backward (blue curve). These effects can be traced to the weights matrices, as can be seen in (B)-(D). **(B)** The mean weights matrices with dominant-forward (corresponding to the green line in (A)). The mean is computed over the active units coding for a memory. **(C)** Same as (B) but for dominant-backward learning rates (corresponding to the blue line in (A)) and **(D)** no heteroassociative component in the learning rule (corresponding to the red line in (A)).

**Figure S6:**
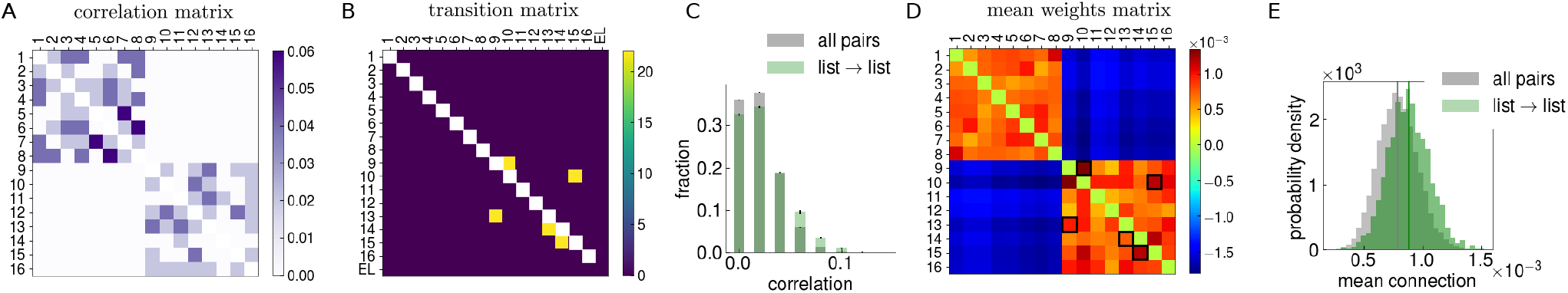
Transitions between memories in the two-set model are guided by stronger connections. **(A)** We study our model with the storage of two sets of memories. Each set contains randomly correlated memories, but the two sets are orthogonal to one another. Pairs of memories belonging to the same set are randomly correlated but pairs in which memories belong to different sets are orthogonal to one another. The matrix expresses the correlation between the memories. The colourbar corresponds to the strength of the correlation. **(B)** The transition matrix for one sample recall trial. The network’s dynamics is entirely restricted to the second set (lower right block), as the network was cued with a memory item in this set. The colourbar corresponds to the number of transitions that have occurred. **(C)** The recall transitions are weakly guided by the correlations in this case. The gray distribution corresponds to the pairwise correlation values between all memories in the first set, while the green distribution corresponds to the correlation values between pairs between which at least one transition has occurred. **(D)** The mean weights between memory items, computed across the different neurons coding for them. In computing this mean, we exclude the neurons that are common to pairs of memory items. The colourbar marks the strength of this mean weight. The black squares mark those pairs of memory items among which transitions occur, as shown in (B). **(E)** The transitions are also guided by the mean connections between memories. The distribution of the mean weights as computed in (D). In gray is the mean weights distribution between all pairs of within-set memories (block diagonal in (D)), whereas in green is the distribution of the mean weights corresponding to pairs of memories between which at least one transition has occurred (black squares in (D)). Panels (A) and (B) also appear in Fig. 6 in the main text and are repeated here for clarity.

## Footnotes

1 For the ON period (after ON switch): 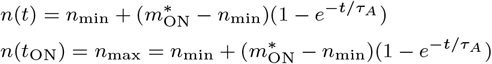 For the OFF period (after OFF switch): 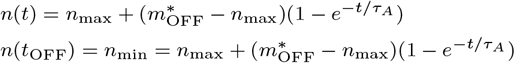

